# LncRNA Malat1 represses Th17 effector program by maintaining a critical bivalent super-enhancer and promotes intestinal inflammation

**DOI:** 10.1101/2022.03.21.485192

**Authors:** Shengyun Ma, Bing Zhou, Yohei Abe, Nicholas Chen, Claire Luo, Anna Zheng, Yuxin Li, Parth R. Patel, Shefali A. Patel, Yajing Hao, John T. Chang, Xiang-Dong Fu, Wendy Jia Men Huang

## Abstract

Interleukin IL-17 cytokines are central regulators of mucosal homeostasis and disease. In mouse models of colonic tissue injury, IL-17A promotes epithelial barrier functions and restricts local inflammation. Here, we report that IL-17A production by the diverse T lymphocyte subsets is dynamically regulated at different stages of colitis pathogenesis. During the onset and peak of the disease, Tγδ17 cells are the major IL-17A producers, while Th17 activity is temporally restricted by long non-coding RNA (lncRNA) Malat1. In response to IL-6 and TGFβ signaling, Malat1 is recruited to the Th17-specific cis-regulatory elements, CNS3 and CNS4, of the *Il17a* locus to fine-tune bivalent super-enhancer activities and repress local transcription. During the resolution phase of inflammation, Malat1 expression is down-regulated to enhance Th17 activities, allowing Th17 cells to emerge as the main producers of IL-17A in the colonic lamina propria. Genetic ablation of Malat1 potentiates IL-17A production in Th17 cells and improves disease outcomes in mouse models of colitis. These findings uncover a surprising role of a chromatin-associated lncRNA in regulating colonic Th17-specific responses to control the timing of inflammation resolution.

**Significance Statement:** T cells are critical modulators of mucosal barrier function and inflammation. The function of long-noncoding RNAs (lncRNAs) in T cells and their role in mucosal inflammation remain elusive. Here, we identify an essential role of the lncRNA Malat1 restricting transcription of the *Il17a* locus in Th17 cells encoding a cytokine implicated in epithelial barrier function post-injury. By controlling the activity of the bivalent super-enhancer at the *Il17a* locus, Malat1 regulates the timing of inflammation resolution in the intestine. The Malat1-*Il17a* pathway reveals new targets for combating mucosal diseases.

**Graphic Abstract:** 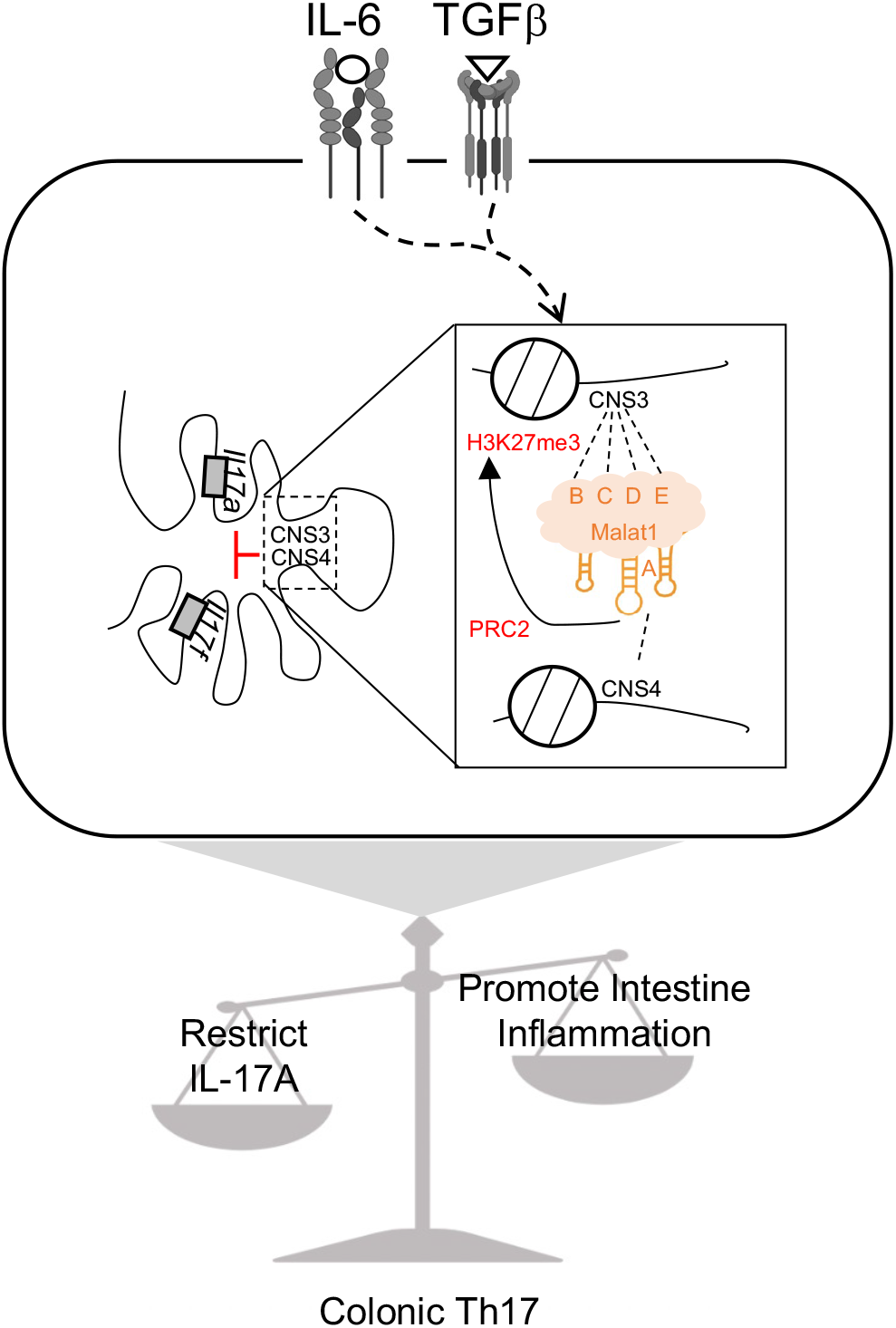

## Introduction

Interleukin 17 (IL-17) maintains intestinal barrier function by stimulating epithelial cell proliferation, tight junction protein expression (1–3), and recruiting myeloid suppressor cells to restrict local inflammation (4). Neutralization of IL-17A or its receptor worsens intestine inflammation in Inflammatory bowel disease (IBD) patients (5, 6). The protective role of IL-17A in the intestine is also observed across multiple mouse models of colitis, including the CD4^+^ T cell transfer (1, 7), the multidrug resistance-1a-ablated (*Abcb1a^-/-^*) (8), and the dextran sulfate sodium (DSS)-induced models (1, 9–11).

IL-17A is secreted by multiple intestinal T cell subsets in a context-dependent and temporally regulated fashion. In the dextran sulfate sodium (DSS) induced colonic injury model, Tγδ17 cells are the primary IL-17A-secreting population at the onset of disease. In the resolution phase, however, Th17 cells surpass Tγδ17 cells as the main source of local IL-17A (1, 4). But how different subsets are regulated and contribute to disease pathogenesis remains poorly defined. Understanding the cell-specific mechanism(s) responsible for the diverse intestinal T cell responses may provide new targets for therapeutic intervention for treating intestinal inflammatory diseases.

Transcription of *Il17a* and its neighboring *Il17f* is controlled by a bivalent super-enhancer (SE) in Th17 cells (12–15). Bivalent SEs are large clusters of enhancers enriched with both active H3K27 acetylation (H3K27ac) and repressive tri-methylation (H3K27me3) marks (16, 17). A bivalent chromatin state is thought to assure the silencing of developmentally regulated genes while keeping them poised for rapid activation during cell differentiation (18, 19). In Th17 cells, multiple transcription factors, including signal transducer and activator of transcription 3 (STAT3), RAR-related orphan receptor gamma (RORγt), SMAD Family Member 2 (SMAD2), and tripartite motif containing 28 (TRIM28), are required for transcriptionally activating the *Il17a-Il17f* locus (12, 20–22), but the molecules and mechanisms that maintain the *Il17a-Il17f* bivalent SE remain elusive.

Emerging evidence indicates that chromatin-associated long non-coding RNAs (lncRNAs) can modulate gene transcription (23, 24). Malat1 is the most abundantly expressed lncRNA in mammalian genomes and is ubiquitously expressed in all cell and tissue types, including T cells (25, 26). Surprisingly, mouse genetic studies show that this lncRNA is dispensable for organ development, cell viability, fertility, and lifespan (27–29). Malat1 is often found on the chromatin of actively transcribed genes, suggesting its vital role in organizing transcriptional hubs in the nucleus (30, 31). Additionally, this lncRNA has also been reported to partner with components of the PRC2 complex to promote histone H3K27 hypermethylation (32–36). However, its involvement in IBD remains controversial, as MALAT1 expression is reported to be down-regulated in Crohn’s disease patients (37), but up-regulated in ulcerative colitis cohorts (38). A recent study utilized the *Malat1^LacZpA/LacZpA (RIKEN)^* knockout mouse line to investigate the function of this lncRNA in the intestine and its contribution to intestinal diseases. However, the interpretation of the reported findings is unclear as *Malat1* in this model is only partially ablated, and results are further confounded by the reduced expression of a *Malat1* nearby gene *Neat1* (28), another abundant lncRNA previously reported to be involved in intestine inflammation (39).

Here, we uncover a surprising role of Malat1 in repressing IL-17 cytokine production in Th17 cells using a well-characterized Malat1 knockout (*Malat1^-/-^*) mouse line in which the Malat1 transcript is completely ablated without detectable effect on Neat1 expression (29). During the resolution phase of colitis, Malat1 is down-regulated, licensing Th17 to surpass Tγδ17 as the major IL-17A producer. Malat1 knockout mice are equipped with elevated IL-17A production capacity in their Th17 cells and are better protected when challenged with DSS-induced colitis. Mechanistically, Malat1 occupies the Th17-specific cis-regulatory element (CNS3 and CNS4) on the *Il17a* locus, recruiting the Polycomb Repressive Complex 2 (PRC2) to restrict the transcriptional activities of the local bivalent super-enhancer.

## Results

### Malat1 expression is dynamically regulated in CD4^+^ T cells during development and in disease settings

In the thymus, Malat1 is modestly expressed in developing T cells of the double negative (DN) to the immature single positive (ISP) stages and is further upregulated in the double-positive (DP) and mature CD4^+^ thymocytes (SP4) (Fig. 1A) (40). Elevated levels of Malat1 are also maintained by naïve CD4^+^, regulatory T cells, and Tγδ cells in the periphery. Notably, *in vitro* activation of naïve CD4^+^ cells resulted in the downregulation of Malat1 (Fig. 1A). Therefore, we first assessed the role of Malat1 in T cell development. Malat1^-/-^ and Malat1^+/+^ (referred to as CTL from now) cohoused littermates were obtained from heterozygote crosses (29). Under steady-state, Malat1^-/-^ and CTL mice had similar body weights (Fig. S1A-B). Thymic T cell populations in the CTL and Malat1^-/-^ mice were comparable (Fig. S1C-D). Steady-state spleens also harbored similar proportions of mature CD4^+^ helper T cells, CD8^+^ cytotoxic T cells, and Treg (CD4^+^CD25^+^) cells between CTL and Malat1^-/-^ mice (Fig. S1E-F). Among the splenic CD4^+^ helper T cells, the ratio of naïve (CD62L^+^CD44^-^) and activated (CD62L^-^CD44^+^) cells was comparable in CTL and Malat1^-/-^ mice (Fig. S1G). These data showed that Malat1 was dispensable for thymic T cell development and mature T cell populations in the secondary lymphoid organ.

**Figure 1.**
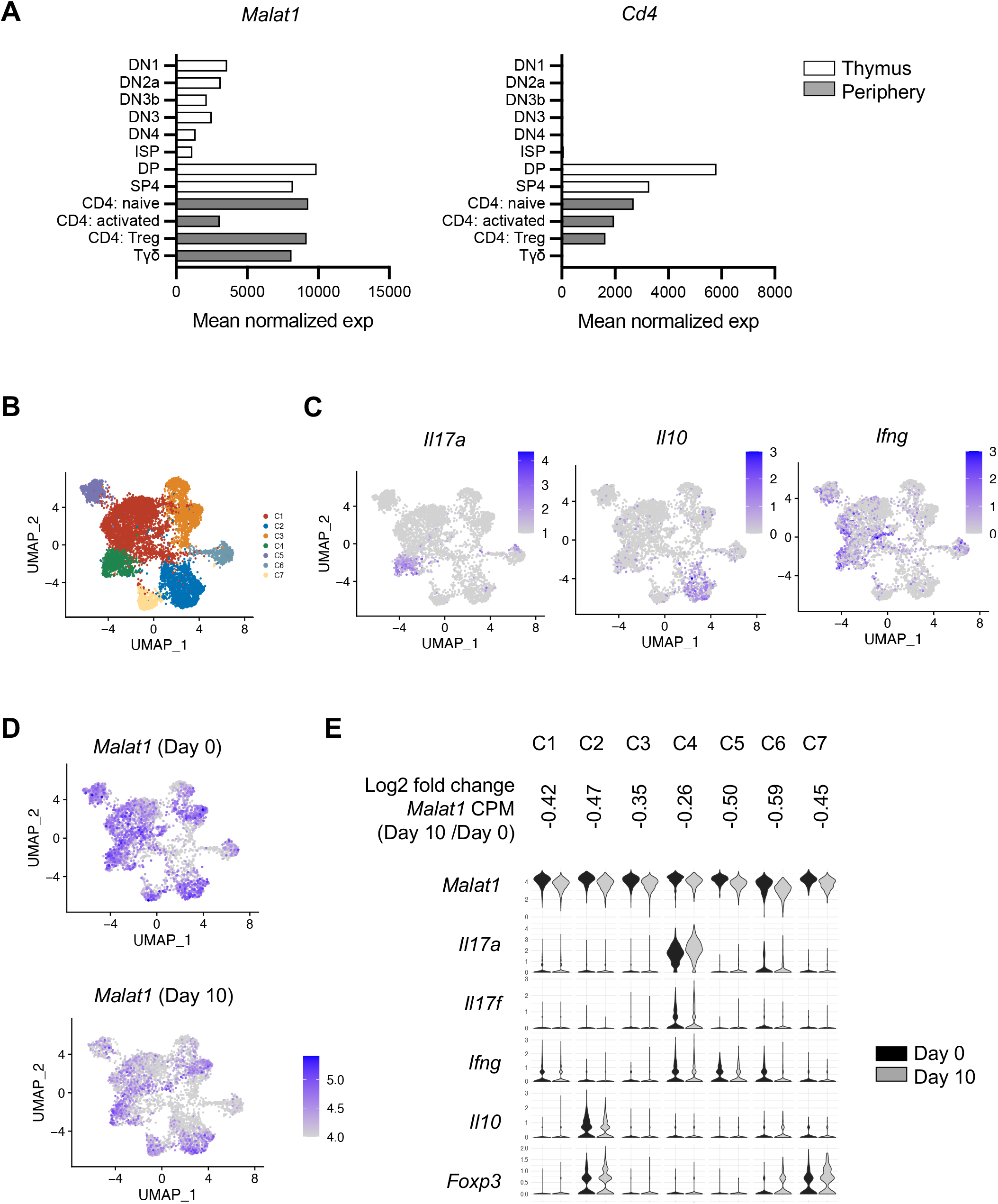
Malat1 expression in CD4^+^ T cells is dynamically regulated during development and in disease settings. A. Expression level of *Malat1* and *Cd4* in different T cell development stages from publicly available RNA-seq (GSE109125). B. UMAP plot of the seven clusters of CD4^+^ T cells, including Th1 (cluster C1), Treg (C2), Naïve (C3), Th17 (C4), Cytotoxic (CTL, C5), proliferating (C6) and Gata3^+^ Treg (C7) cells from colonic lamina propria of CTL (DSS day 0) and DSS treated mice (DSS day 10). C. UMAP plots of *Il17a, Il10*, and *Ifng* expressions in cLP CD4^+^ T cells. D. UMAP plots of *Malat1* and *Il17a* expressions in cLP CD4^+^ T of CTL and DSS treated mice. E. Violin plot of indicated gene expressions in different T cell subsets of CTL and DSS treated mice.

Next, we assessed the level of Malat1 in mucosal resident CD4^+^ T cells under steady-state and asked whether its expression is perturbed upon colonic injury. Single-cell RNA-seq analysis of the colonic lamina propria CD4^+^ T cells from mice challenged with or without Dextran Sulfate Sodium (DSS), a model of colitis where recent studies have implicated the role of CD4^+^ T cells in contributing to disease severity (41–46), showed that Malat1 was abundantly expressed in different subsets of colonic T cells under steady-state (Fig. 1B-C and S2A) (47) and its expression was downregulated during the inflammation resolution phase of DSS-induced colitis (Fig. 1D-E and Table S1).

### Malat1 promotes colitis and regulates colonic Th17 function under inflamed settings

Based on the above observations, we asked whether Malat1 contributed to the differentiation and function of colonic resident T cells under homeostatic and inflamed settings. Using the gating strategy outlined in Fig S3A, we examined the resident T cells in the steady-state colonic lamina propria in the CTL and Malat1^-/-^ mice, but did not detect a change in the proportions and cell numbers of Th17 cells (defined as CD4^+^RORγt^+^Foxp3^-^), RORγt^+^ Treg (defined as CD4^+^RORγt^+^Foxp3^+^), conventional Tregs (cTreg, defined as CD4^+^RORγt^-^Foxp3^+^), Tγδ17 (defined as CD4^-^TCRγδ^+^RORγt^+^), and ILC3s (defined as CD4^-^TCRγδ^-^RORγt^+^) (Fig. S4A-B), suggesting that Malat1 is not involved in T cell differentiation and their residency in the colonic tissue under steadystate. Furthermore, the proportions of IL-17A expressing colonic Th17 cells in the Malat1^-/-^ mice were similar to those found in CTL mice under steady state (Fig. S4C-D). To investigate the role of Malat1 in settings of inflammation, cohoused Malat1^-/-^ and CTL littermates were challenged with 2% DSS for seven days. On day 8-10, Malat1^-/-^ mice experienced less weight loss compared to CTL littermates (Fig. 2A). Colonic tissues from Malat1^-/-^ mice displayed significantly less muscle thickening and submucosal inflammation (Fig. 2B-C). These data revealed that Malat1 is a positive regulator of intestine inflammation *in vivo*.

**Figure 2.**
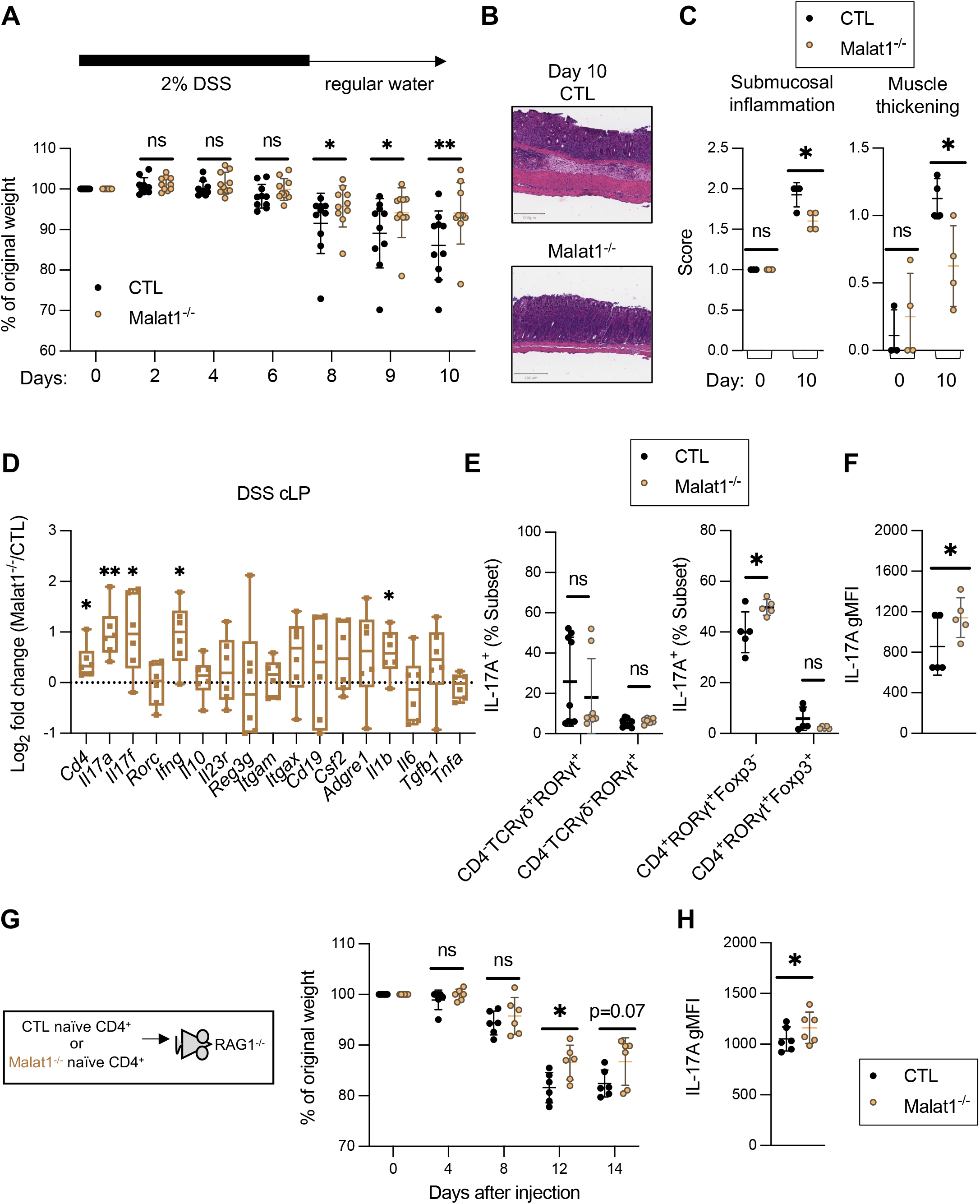
Malat1 in CD4^+^ T helper cells augments colonic inflammation. A. Weight change of CTL (Malat1^+/+^ or Malat1^+/-^) (black, n=10) and Malat1^-/-^ (red, n=9) cohoused littermates challenged with 2% DSS in drinking water for 7 days and regular water for an additional 3 days. Results were combined from three independent experiments. Each dot represents result from one mouse. * p-value<0.05, ** p-value<0.01, ns: not significant (multiple t-test). B. Representative H&E staining of colonic sections from CTL and Malat1^-/-^ mice harvested on DSS day 10. Scale bar: 200 μm. C. Colonic submucosal inflammation and muscle thickening scores in CTL (n=4) and Malat1^-/-^ (n=4) mice treated with or without DSS. DSS-treated tissues were harvested on day 10. Each dot represents result from one mouse. * p-value<0.05, ns: not significant (t-test). D. Summarized log_2_ fold change of relative RNA levels of the indicated genes in total colonic lamina propria mononuclear cells harvested from CTL (n=6) and Malat1^-/-^ (n=6) mice on day 10 post initial DSS treatment as detected by qRT-PCR. Each dot represents result from one mouse. * p-value<0.05, ** p-value<0.01, ns: not significant (multiple t-test). E. The proportion of IL-17A^+^ cells among the different immune cell subsets from CTL (n=5) and Malat1^-/-^ (n=5) littermates harvested on day 10 post DSS treatment. Each dot represents result from one mouse. * p-value<0.05, ns: not significant (multiple paired t-test). F. IL-17A gMFI in cLP Th17 cells (defined as CD4^+^ RORγt^+^Foxp3^-^) from F. * p-value<0.05, (paired t-test). G. Weight changes of RAG1^-/-^ mice receiving CTL or Malat1^-/-^ naive CD4^+^ T cells. Each dot represents result from one RAG1^-/-^ mouse. * p-value<0.05 (multiple t-test, n=6). H. IL-17A gMFI in cLP Th17 cells (defined as CD4^+^ RORγt^+^Foxp3^-^) from G. * p-value<0.05, (paired t-test).

In total colonic lamina propria (cLP) cell lysates from DSS-challenged Malat1^-/-^ mice, the transcripts encoding IL-17A were significantly elevated at the RNA level (Fig. 2D and S5A). This difference was not due to a change in the number of Th17, RORγt^+^ Treg, Tγδ17, or ILC3 (Fig. S5B), suggesting that Malat1 is not involved in the differentiation and/or maintenance of these populations. For other T cell populations previously implicated in colitis, such as conventional Tregs and Th1, their proportions, cell numbers, and cytokine production potential were also similar between DSS-challenged CTL and Malat1^-/-^ mice (Fig. S5D-E, representative results in Fig. S6A). However, the proportion and number of IL-17A-expressing Th17s were higher in the DSS-challenged Malat1^-/-^ colons (Fig. 2E, representative results in Fig. S6A). The geometric mean fluorescent intensity (gMFI) of individual IL-17A-expressing Malat1^-/-^ Th17 cells was also higher (Fig. 2F), suggesting that Malat1^-/-^ Th17 cells had higher cytokine production potential on a per-cell basis in the DSS-challenged colon. These results demonstrate that Malat1 negatively regulates Th17 cytokine production capacities in a context-dependent and subset-specific manner.

Similar to the DSS colitis model, *Il17a* transcript levels also negatively correlated with *Malat1* abundance in a chronic colitis model induced by the transfer of naive T cells into RAG-deficient mice (Fig. S7A) (48). To test the role of Malat1 in this model, we isolated naïve CD4^+^ T cells, confirmed to have >98% purity by flow cytometry (Fig. S7B), from CTL and Malat1^-/-^ littermates and transferred them to RAG1^-/-^ recipients. Consistent with our observation in the DSS model, RAG1^-/-^ mice transplanted with Malat1^-/-^ CD4^+^ T cells experienced less weight loss compared to those receiving CTL cells (Fig. 2G), and the signals of IL-17A in colonic Malat1^-/-^ Th17 cells were also higher (Fig. 2H). Together, these data demonstrated that Malat1 in CD4^+^ T helper cells is sufficient to drive disease pathogenesis in models of colitis.

### Malat1 regulates Th17 function in a cell-autonomous manner

To determine the cell-intrinsic role of Malat1 in regulating Th17 differentiation and function, we isolated naïve T cells from CTL and Malat1^-/-^ mice (Fig. S7B) and differentiated these cells toward Th17 cells in the presence of IL-6 and TGFβ. Malat1 ablation augmented the proportions of IL-17A and IL-17F (Fig. 3A), similar to our *in vivo* observations and consistent with results from a previous study using a transient knockdown approach (49). Importantly, the augmented cytokine levels observed in Malat1^-/-^ Th17 cells were not due to altered expression of RORγt, the master transcription factor for the genes encoding these two cytokines (Fig. 3A and S8A-C). On a per-cell basis, the gMFI of IL-17A and IL-17F were significantly higher in the Malat1^-/-^ Th17 cells (Fig. 3B). At the transcript level, there was also a significant elevation of *Il17a* and *Il17f* abundance in cultured Malat1^-/-^ Th17 cells (Fig. 3C), suggesting that Malat1 likely regulates IL-17A and IL-17F at the chromatin and/or RNA levels. In contrast, Malat1 was dispensable for *in vitro* differentiation of naïve T cells toward Treg, Th1, and Th2 lineages (Fig. S9A-D). These findings established a selective role of Malat1 in regulating Th17 cell effector function in a cell-intrinsic manner.

**Figure 3.**
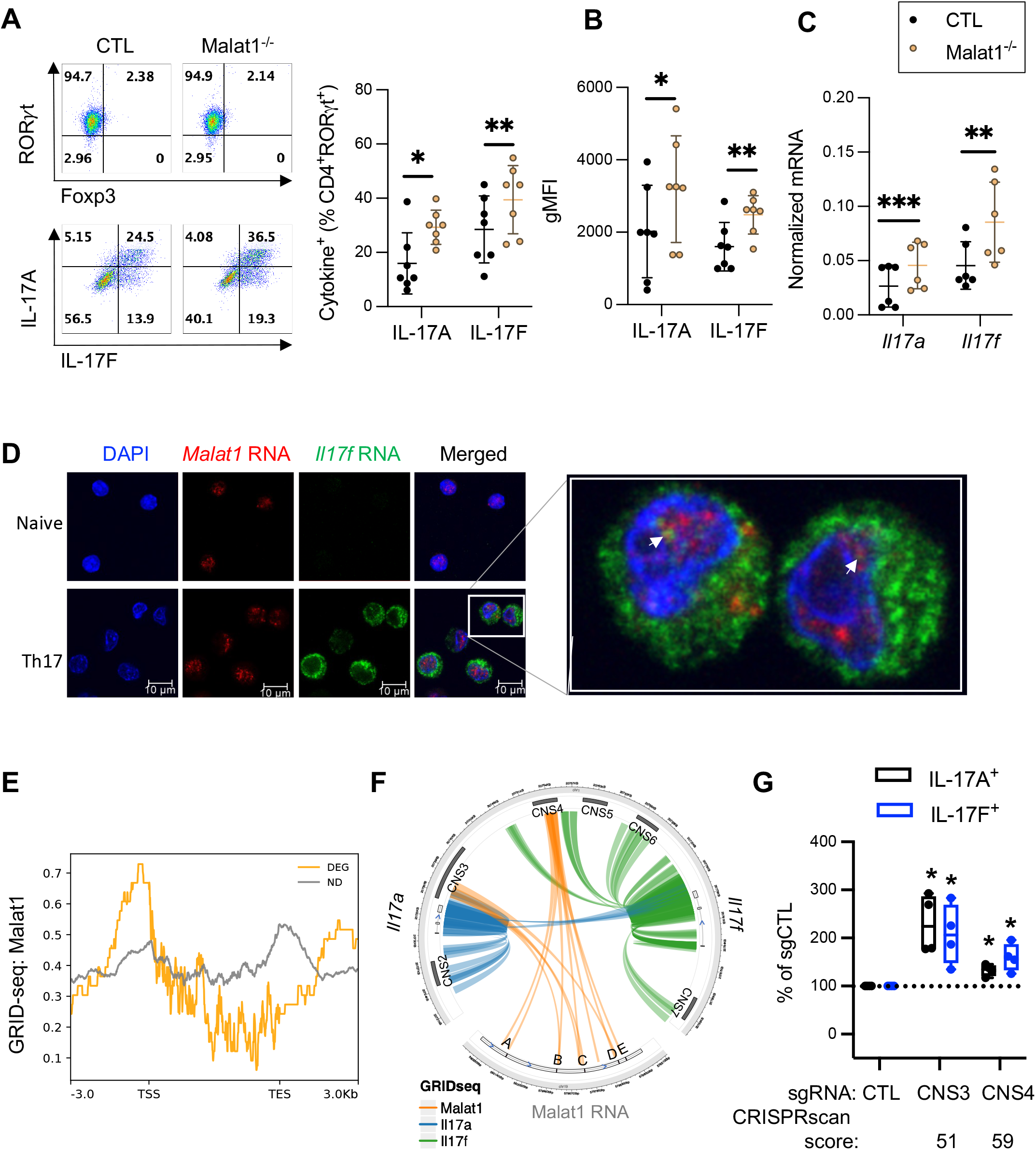
Malat1 occupies select regulatory elements at the Il17a-Il17f super-enhancer in cultured Th17. A. Left: representative flow cytometry analysis of RORγt, Foxp3, IL-17A, and IL-17F expression of CTL and Malat1^-/-^ Th17 cells cultured with IL-6 and TGFβ for 3 days. Right: Summarized proportion of IL-17A and IL-17F in CTL and Malat1^-/-^ RORγt^+^ Th17 cells cultured with IL-6 and TGFβ for 3 days. Each dot represents result from one independent culture. * p-value<0.05, ** p-value<0.01 (n=8, multiple paired t-test). B. Summarized gMFI of IL-17A and IL17-F in Th17 cells cultured with IL-6 and TGFβ for 3 days. * p-value<0.05, ** p-value<0.01 (multiple t-test, n=7). C. Normalized mRNA of *Il17a* and *Il17f* relative to *Gapdh* in cultured CTL and Malat1^-/-^ Th17 cells as detected by qRT-PCR. Each dot represents result from one independent culture. ** p-value<0.01, *** p-value<0.001 (multiple paired t-test, n=6). D. Immunofluorescence *in situ* hybridization (RNA-FISH) detection of Malat1 RNA (red) and *Il17f* (green) mRNA in cultured CTL naïve and Th17 cells. DAPI (blue): nuclear DNA stain. White arrows: regions of *Malat1* and *Il17f* RNA co-localization in the Th17 nucleus. Scale bar: 5 μm. E. Malat1 chromatin occupancy on the gene body of Malat1 dependent genes (orange line, DEG with a log_2_ fold change cutoff of <-0.4 or >0.4) and 1000 randomly selected Malat1 independent Th17 genes (black line, ND) as determined by GRID-seq. F. Circos plot showing the interactions between RNAs transcribed from *Il17a* (blue), *Il17f* (green), *Malat1* (orange) loci and chromatin DNA at the *Il17a-Il17f* locus in CTL Th17 cells captured by GRID-seq. Five regions on the Malat1 RNA found in close proximity to the *Il17a-Il17f* locus were designated as A-E. G. Summarized log_2_ fold change of flow cytometry analysis of IL-17A and IL-17F in Cas9^+^ Th17 cells transduced with retroviral constructs expressing sgRNAs targeting CNS3 and CNS4 noted in C relative to those with CTL (empty vector). CRISPRscan scores were calculated according to (84). The higher the score predicts greater Cas9 cleavage efficiency. Each dot represents result from one independent culture. * p-value<0.05 (t-test, n=4).

### Malat1-dependent gene expression in Th17 cells

Then we asked whether Malat1 regulates the expression of other genes beyond *Il17a* and *Il17f* in Th17 cells polarized in the presence of IL-6 and TGFβ. RNA-seq analysis confirmed an increase of *Il17a* and *Il17f* transcripts in cultured Malat1^-/-^ Th17 cells (Fig. S10A-B) and identified additional Malat1-regulated genes encoding molecules involved in Th17 cell differentiation (*Pp3ca*), cell migration (*Arhgap5, Itgb1*), senescence (*Zfp36l1, Zfp36l2*), Hippo signaling (*Bmpr2, Ajuba*), and metabolism (*Dgat1, Aldh6a1)* (Fig. S10C). Surprisingly, most of the Malat1-dependent genes resided outside of the Malat1 resident chromosome 19 (Fig. S10D), suggesting that Malat1 has regulatory footprints on distal chromatins in Th17 cells. qRT-PCR analysis on cultured Th17 cells generated from independent pairs of CTL and Malat1^-/-^ littermates confirmed several additional Malat1-regulated genes (Fig. S10E), extending the regulatory role of Malat1 beyond cytokine production.

### Genome-wide Malat1 localization in Th17 cells

In mouse embryonic fibroblasts and interstitial cells of testis, Malat1 preferentially resides in the nuclear compartment and enriches on the active chromatins (29, 50). Therefore, we asked whether Malat1 also localizes to the nucleus in T cells. Single-molecule RNA FISH assay revealed that the majority of Malat1 indeed localized to the nucleus in both naïve and cultured Th17 cells (Fig. 3D). Interestingly, a subset of Malat1 RNAs colocalized with the *Il17f* transcription hubs (indicated by triangles in Fig. 3D). Therefore, we hypothesized that Malat1 might be recruited to the chromatin at or near the vicinity of the *Il17a-Il17f* locus to regulate local transcription.

To unbiasedly characterize the chromatin occupancy of Malat1 in Th17 cells, we used the GRID-seq approach recently developed to capture RNA-DNA interactions in the native nuclear 3D space (51). From two independent replicates of wildtype Th17 cells, we identified a total of 7,039 enriched Malat1-chromatin interactions (peaks), mainly on autosomes (Fig. S11A). Unlike previous reports (30, 52), the majority of Malat1 peaks in Th17 cells (54.5%) were on gene promoters. 35.2% and 10.3% were found on gene bodies and intergenic regions, respectively (Fig. S11B and Table S2). Malat1 preferentially occupied active regulatory elements (defined as open chromatin sites enriched with H3K27ac signals). 91.9% of the Malat1 occupied promoters harbored open chromatin and 75.3% of the Malat1 occupied intergenic regions harbored open chromatin. The remaining one-quarter of the Malat1 peaks found on intergenic regions with closed chromatin likely indicate a role of Malat1 in heterochromatin and gene repression in line with previous studies (32–36, 53). On Malat1-dependent Th17 genes (DEG), Malat1 is preferentially enriched near the transcriptional start site (TSS) (Fig 3E). These results extend the role of Malal1 in organizing active gene hubs (30, 31, 52) to having bivalent roles in directing both gene activation and repression in Th17 cells.

### Malat1 contributes to the recruitment of PRC2 to the *Il17a-Il17f* bivalent super-enhancer

In naïve cells, the *Il17a-Il17f* locus displayed a closed chromatin configuration with little detectable Malat1 binding (Fig. S12A-B). In cultured Th17 cells, Malat1 occupied two open elements within the previously defined cis non-coding site CNS3 and CNS4 (54) (Fig. 3F and S12A-B). Previous studies also reported that CNS3 and CNS4 harbor open chromatin configuration in intestinal Th17 cells (Fig. S12C) (55). Interestingly, both CNS3 and CNS4 in Tγδ17 cells were kept in a closed chromatin configuration (Fig. S12C) (56). These subset-specific chromatin landscape patterns correlate with the ability of Tγδ17 cells to evade Malat1 regulation of IL-17 (Fig. 2E and S4C-D). When CNS3 or CNS4 was targeted by Cas9-sgRNA mediated mutagenesis in wildtype Th17 cells, IL-17A and IL-17F productions were significantly upregulated (Fig. 3G), suggesting that these Malat1 bound sites serve as repressive regulatory elements for the *Il17a-Il17f* locus.

Furthermore, the chromatin region spanning from CNS2 to CNS6 had the characteristics of a bivalent super-enhancer, decorated by both active and repressive marks including H3K27ac and H3K27me3 (Fig. 4A). H3K27me3 deposition can be mediated by the PRC2 complex (57, 58) and members of this complex, including EZH2 and SUZ12, have been recently reported to bind Malat1 (35, 59, 60). On the *Il17a-Il17f* super-enhancer, EZH2 and SUZ12 signals clustered around CNS3, CNS4, and CNS5 (Fig. 4A). Therefore, we tested whether Malat1 is involved in the recruitment of PRC2 and/or H3K27me3 deposition on the *Il17a-Il17f* bivalent SE to modulate the transcriptional output of the locus. ChIP-qPCR analysis of CTL and Malat1^-/-^ Th17 cells indeed revealed that Malat1 ablation selectively and significantly diminished SUZ12 binding at CNS3 and CNS4 (Fig. 4B). ChIP-seq also revealed modest Malat1-dependency of H3K27me3 signals at CNS3 and CNS4 in Th17 cells (Fig. 4C). No change was observed at other regions of the *Il17a-Il17f* SE, such as CNS5, and H3K27 acetylation between CTL and Malat1^-/-^ Th17 cells were comparable (Fig. 4D). Together, these results demonstrate that Malat1 facilitates the recruitment of PRC2 to maintain the SE bivalency of the *Il17a-Il17f* locus (modeled in Fig. 4E).

**Figure 4.**
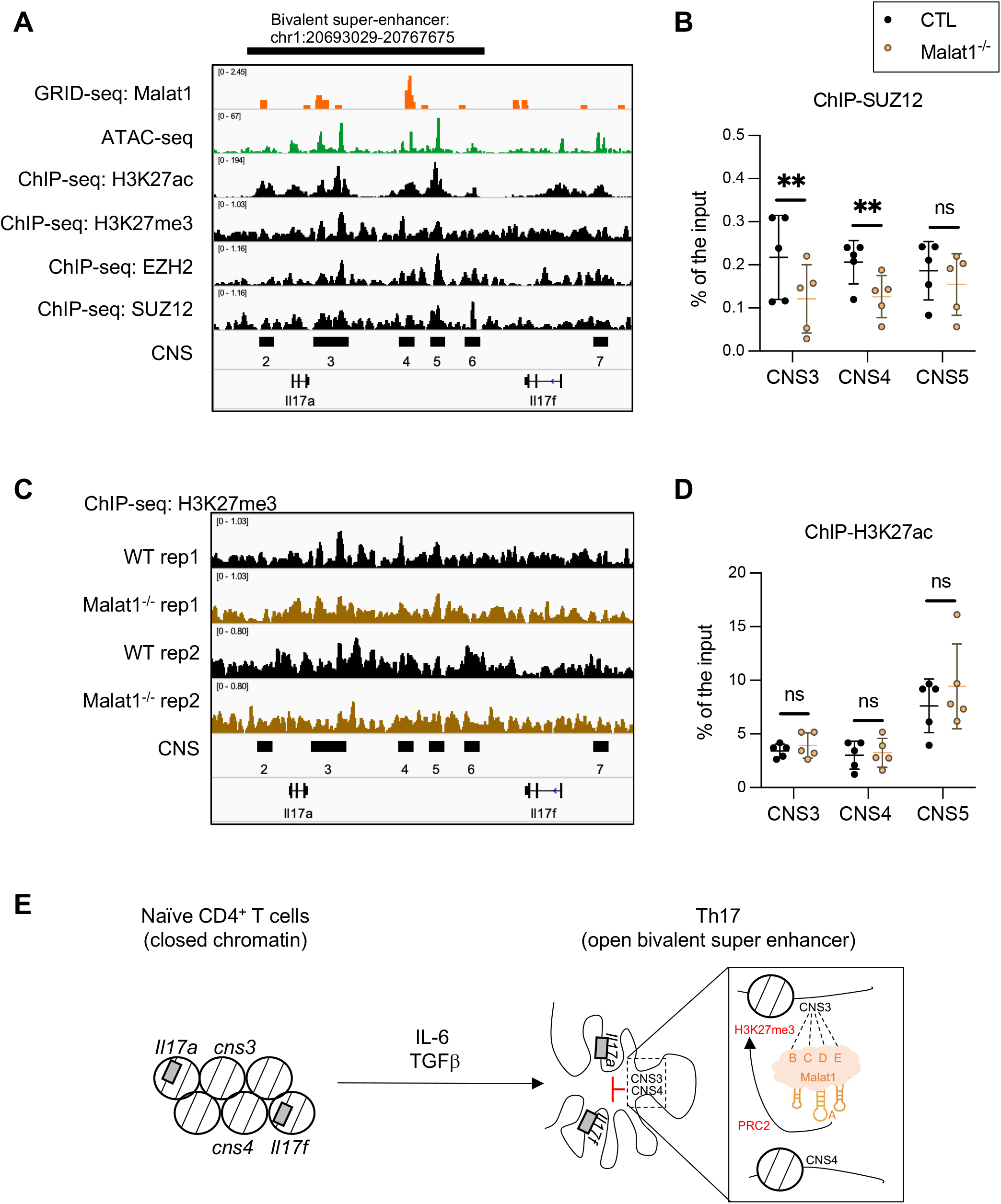
Malat1 recruits SUZ12 to modulate *Il17a-Il17f* SE bivalency in Th17 cells. A. IGV browser view of Malat1, H3K27ac, H3K27me3, EZH2, and SUZ12 signals on the *Il17a-Il17f* locus in Th17 cells as detected by GRID-seq and ChIP-seq. B. ChIP-qRT analysis of SUZ12 at the *Il17a-Il17f* locus in cultured Th17 cells from CTL (n=5) and Malat1^-/-^ (n=5) littermates. Each dot represents result from one independent culture. p, promoter. Malat1 occupied sites (orange shade). ** p-value<0.01, ns: not significant (multiple paired t-test). C. IGV browser view of H3K27me3 signals on the *Il17a-Il17f* locus in CTL (n=2) and Malat1^-/-^ (n=2) Th17 cells as detected by ChIP-seq. D. ChIP-qRT analysis of H3K27ac at the *Il17a-Il17f* locus in cultured Th17 cells from CTL (n=5) and Malat1^-/-^ (n=5) littermates. Each dot represents result from one independent culture. p, promoter. Malat1 occupied sites (orange shade). ns: not significant (multiple paired t-test). E. Model: upon IL-6 and TGFβ signaling, nucleosome remodeling opens up CNS2 to CNS6 on the *Il17a-Il17f* locus and chromatin looping ensures these distal elements are kept in close spatial proximity. Subsequently, CNS3 and CNS4 on the *Il17a-Il17f* locus serve as a physical dock for Malat1 recruitment. Notably, regions B-E on Malat1 provide multiple and redundant contact points for anchoring to CNS3 and region A on Malat1 is uniquely oriented in close spatial proximity to CNS4. Together, Malat1 facilitates the recruitment of SUZ12 and H3K27me3 deposition to the *Il17a-Il17f* locus.

On two other confirmed Malat1-repressed targets, *Dgat1* and *Tecpr1* (Fig. S10E), Malat1 was found on their promoters, intragenic, and distal regulatory elements (Fig. S13A). In the Malat1^-/-^ Th17 cells, SUZ12 or H3K27me3 enrichment on the *Dgat1* and *Tecpr1* promoters was significantly reduced (Fig. S13B). These results suggest that the partnership of Malat1 and PRC2 we observed on the *Il17a-Il17f* super-enhancer also applies to other Th17 targets.

### Identifying the Malat1 RNA region necessary for repressing IL-17 in Th17 cells

On the Malat1 transcript, five regions made close contact with the *Il17a-Il17f* chromatin (Fig. 3F, referred to as regions A to E). Three of the five regions (B, C, and D) also made frequent contact with other Malat1 targets (Fig. 5A). These results suggest that both shared and gene-specific mechanisms likely underly Malat1 regulation of Th17 targets and that regions A and E may serve a unique role on the *Il17a-Il17f* locus. To test this possibility, we used CRISPR technology to introduce indel mutations in different regions of Malat1. Briefly, we transduced activated CD4^+^ T cells from Cas9 transgenic mice with retroviruses carrying region-specific sgRNA expression constructs. After confirming Cas9-sgRNA-induced indel mutations by Sanger sequencing, we assessed whether specific sgRNA(s) alter the expression of IL-17A and IL-17F. Interestingly, we found two independent sgRNAs, sgA.1 and sgA.2 targeting region A (chr19: 5801450-5801521, GRCm38/mm10), but not sgRNAs targeting other Malat1 regions (B-E), significantly augmented IL-17A and IL-17F at the mRNA and protein levels (Fig. 5B-D), demonstrating that region A is necessary for Malat1-mediated repression of cytokine production in Th17 cells. Interestingly, region A on Malat1 was dispensable for regulating other direct target genes, such as *Syncrip, Dgat1*, and *Ddx3x* (Fig. 5D). Importantly, targeting region A did not alter the overall Malat1 RNA abundance (Fig. 5D), indicating that the impact on IL-17A and IL-17F is likely due to a change in Malat1 RNA structure and/or function.

**Figure 5.**
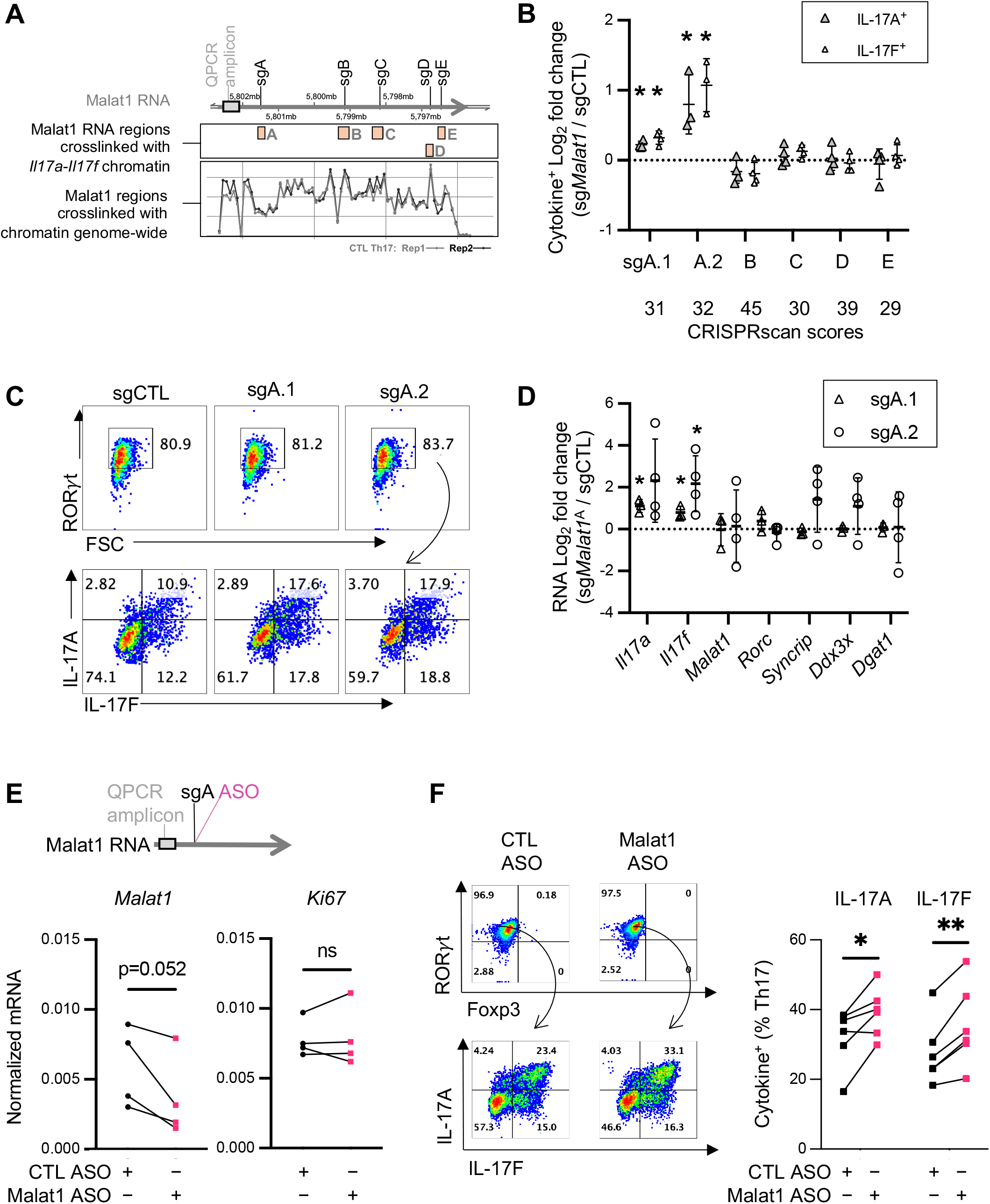
A Malat1 RNA region implicated in *Il17a-Il17f* transcriptional repression. A. Comparing Malat1 RNA regions enriched on Th17 chromatin genome-wide and those found in close proximity to *Il17a-Il17f* chromatin (orange blocks) from 4C. Vertical black lines indicate positions of small guide (sg) RNAs designed to mutate different regions of Malat1. B. Summarized log_2_ fold change of IL-17A and IL-17F-expressing Cas9^+^ Th17 cells transduced with retroviral constructs expressing sgRNAs targeting each *Malat1* regions noted in A relative to those with CTL (empty vector). Each dot represents result from one independent culture. * p-value<0.05 (t-test). CRISPRscan scores were calculated according to (84). The higher the score predicts greater Cas9 cleavage efficiency. C. Representative flow cytometry analysis of RORγt, IL-17A, and IL-17F expression from B. D. Summarized log_2_ fold change of relative RNA levels of the indicated genes in cultured Cas9^+^ Th17 cells transduced with retroviral constructs expressing sgA.1 and sgA.2 relative to those with CTL vector. Each dot represents result from one independent culture. * p-value<0.05 (multiple t-test, n=4). E. Top: position of custom ASO targeting the Malat1 transcript. Bottom: Normalized RNA level of *Malat1* and *Ki67* in ASO-treated cells. Each dot represents result from one mouse. ns: not significant (t-test, n=4). F. Left: representative flow cytometry analysis of RORγt, Foxp3, IL-17A and IL-17F in cultured Th17 cells treated with CTL or Malat1-specific ASO. Right: summarized flow cytometry analysis of IL-17A and IL-17F in cultured Th17 cells treated with CTL or Malat1-specific ASO. Each dot represents result from one independent culture. * p-value<0.05, ** p-value<0.01 (paired t-test, n=5).

Furthermore, we also designed an anti-sense nucleotide (ASO) that complementarily binds to region A. *In vitro* cultured Th17 cells treated with this Malat1-specific ASO showed a modest, but insignificant, reduction of Malat1 RNA and little change of *Ki67*, an index of cell proliferation (Fig. 5E). Malat1-specific ASO treated RORγt-expressing Th17 cells had greater IL-17A and IL-17F production potentials compared to cells treated with CTL ASO (Fig. 5F). Together, these results demonstrates that region A of Malat1 harbors a gene-specific role in repressing cytokine expressions in Th17 cells.

## Discussion

IL-17A is a cytokine important in mucosal barrier maintenance with protective roles during intestinal inflammation (1–3, 8, 9, 11, 61). In the DSS-induced model of colitis, Tγδ17 cells are the primary source of early IL-17A production. As the disease progresses, the protective role of Tγδ17 cells is taken over by the gradually increased IL-17A-expressing Th17 cells emerging as the major source of IL-17A for tissue repair (1, 4). Although both Tγδ17 and Th17 cells are thought to be preprogrammed to express IL-17A, the mechanism(s) restricting early Th17 cell responses and how their effector function gets turned on later during colitis has long been elusive in the field. We now identified the lncRNA Malat1 as a negative regulator of IL-17A expression in colonic Th17 cells post epithelial injury.

Our results demonstrate this regulation occurs on the *Il17a-Il17f* chromatin involving two steps. First, IL-6 and TGFβ signaling facilitate nucleosome remodeling to open up CNS2 to CNS6 on the *Il17a-Il17f* locus and maintain their close physical proximity by chromatin looping. Subsequently, both transcriptional coactivators and corepressors, including Malat1, gain access to their respective sites. Malat1 binding to CNS3 and CNS4 facilitates the recruitment of PRC2 and fine-tunes Th17 cytokine production quantitatively and temporally during colitis (modeled in Fig. 4E). It will be important for future studies to determine which components of the IL-6 and TGFβ signaling machinery, including the SMAD family of transcription factors, are involved in the proper positioning of Malat1 on the Th17 chromatin and in promoting specific Malat1-PRC2 complex assembly to exert a repressive role on the *Il17a-Il17f* locus.

In addition to *Il17a-Il17f*, we also demonstrated that PRC2 occupancy and H3K27 methylation on other Th17 genes, such as *Dgat1*, and *Tecpr1*, also rely on Malat1. These findings in Th17 cells are consistent with our recent report on CD8^+^ T cells (53), providing a new paradigm for a lncRNA to function as the central player in maintaining SE bivalency to regulate T cell functions. It will be also interesting for future studies to examine how enhancer landscape in other IL-17 cytokine expressing immune cell subsets, such as RORγt^+^ Treg, NK17, and MAIT17 cells, influence their sensitivity toward Malat1 regulation. It is important to note that although we found *Il17a-Il17f* in Tγδ17 cells and pathogenic Th17 cells, as well as IL-10 and IFNg production potentials in cTreg and Th1 cells to be Malat1-independent, future transcriptomics studies are needed to decipher whether other aspects of their cellular functions may be regulated by Malat1.

Results of the T cell transfer colitis experiments indicate that Malat1-deficiency in T cells is sufficient to protect against colitis. In the DSS model, however, future studies where Malat1 can be conditionally depleted in select immune cell populations would help better define the extent its expression in T cells versus other cell types contributes to the phenotype observed. We speculate that the elevated colonic IL-17A and/or IL-17F levels from the Malat1^-/-^ Th17 cells post-injury may also alter epithelial cell programs involved in tight junction assembly, epithelial regeneration, and anti-microbial peptide production (2, 3, 8, 61), as well as enhance the recruitment of neutrophils and macrophages for better microbial clearance (62). It remains an open question how these changes alter the overall microbial abundance and/or act upon particular microbial species known to drive Th17-specific immune responses (63–65).

In summary, our results shed new light on the chromatin localization and immunological function of lncRNA Malat1 in T cells, highlighting the previously unrecognized role of this lncRNA in restricting the mucosal protective function of Th17 cells and promoting intestine inflammation.

## Materials and Methods

### Mice

C57BL/6 wild-type (Stock No: 000664), Cas9 (Stock No: 026179), and RAG1^-/-^ (Stock No: 002216) were obtained from the Jackson Laboratory. Malat1^-/-^ mice were obtained from Dr. David Spector’s laboratory and have been previously described (29). Heterozygous mice were bred to yield 6-8 weeks old CTL (Malat1^+/+^ or Malat1^+/-^) and Malat1^-/-^ littermates. Gender-matched cohoused littermates at least eight weeks old were used. All animal studies were approved and followed the Institutional Animal Care and Use Guidelines of the University of California San Diego.

### DSS-induced and T cell transfer colitis

Mice were provided 2% (w/v) DSS (160110; MP, Bio-medicals) in their drinking water for 7 days, followed by 3 days of access to regular drinking water. Mice weights were monitored daily, and tissues were harvested on day 10. For histopathological analysis, distal colons were opened longitudinally, rinsed with phosphate-buffered saline (PBS), and fixed overnight at room temperature in 10% Formalin (Fisher Chemicals). Sections were stained with hematoxylin and eosin (H&E). Images were acquired using the AT2 Aperio Scan Scope. The method for colitis scoring was modified from Koelink et al (66). Three random regions on each colonic tissue were scored in a double-blind fashion and the average of the three scores were graphed for each tissue. The ‘submucosal inflammation’ score (0-3) is defined by the amount of immune cell infiltration in each section (0, none; 1, an increased presence of inflammatory cells; 2, infiltrates also in submucosa; 3, transmural immune infiltrate). The ‘muscle thickening’ score (0-3) is defined by the increase of thickness compared to the standard muscle thickness (300μm) found in wildtype colons in our colony under steady state (0, none; 1, slight; 2, strong; 3, excessive).

For adoptive transfer model of colitis, mouse splenocytes were subjected to ACK lysis followed by anti-CD4 positive selection (Miltenyi Biotec). Naive CD4^+^ T cells (CD4^+^CD25^-^CD44^-^CD62L^+^) were captured by the MA900 Multi-Application Cell Sorter (Sony). 0.5 million sorted naive CD4^+^ T cells were injected intraperitoneally into RAG1^-/-^ recipients. Mice weights were measured twice a week.

### Flow cytometry

To probe for cytokine production potential, cells were stimulated with 5 ng/mL Phorbol 12-myristate 13-acetate (PMA, Millipore Sigma) and 500 ng/mL ionomycin (Millipore Sigma) in the presence of GolgiStop (BD Bioscience) for 5 hours at 37 °C. Antibodies for cell surface molecules were incubated with intact live cells. Following fixation and permeabilization (eBioscience), antibodies for intracellular transcription factors and cytokines were added. Antibodies are listed in Table S3.

### *In vitro* T cell culture

Mouse naive T cells were purified from spleens and lymph nodes of 8-12 weeks old mice using the Naive CD4^+^ T Cell Isolation Kit according to the manufacturer’s instructions (Miltenyi Biotec). Cells were cultured in Iscove’s Modified Dulbecco’s Medium (IMDM, Sigma Aldrich) supplemented with 10% heat-inactivated FBS (Peak Serum), 50 U penicillin-streptomycin (Life Technologies), 2 mM glutamine (Life Technologies), and 50 μM β-mercaptoethanol (Sigma Aldrich). For cell polarization, naive cells were seeded in 24-well or 96-well plates pre-coated with rabbit anti-hamster IgG and cultured for 72 hours in the presence of 0.25 μg/mL anti-CD3ε (eBioscience), 1 μg/mL anti-CD28 (eBioscience) and different cytokines for different cell differentiation. IL-2 (15 U/mL) and IL-12β (10 ng/mL) for Th1; IL-2 (15 U/mL) and IL-4 (10 ng/mL) for Th2; IL-2 (15 U/mL) and TGFβ (5 ng/mL) for Treg; IL-6 (20 ng/mL) and TGFβ (0.1 ng/mL) for Th17. All the cytokines were obtained from R&D Systems.

Anti-sense nucleotide (ASO) experiments were designed following best practice guidelines (67). 2’-deoxy-2’-fluoro-D-arabinonucleic acid (2’-FANA) containing CTL ASOs targeting a Trypanosoma gene (GCCGTTTACGTCGTCACAGA) or Malat1 (TTCTTTGCCTATCTTGAATGC) were purchased from AUM BIOTECH and their purities were confirmed by MALDI. ASOs were incubated with activated CD4^+^ T cell *in vitro* for 48hrs according to manufacturer instructions (AUM BIOTECH).

### Retrovirus transduction in T cells

Malat1-specific sgRNAs were designed by CHOPCHOP (68). sgRNA sequences are listed in Table S4. pSIN retroviral constructs carrying expression cassettes for sgRNA along with a surface protein CD90.1 (Thy1.1) were generated as described previously (69). PlatE cells were used to generate retrovirus as described (70). Virus transduction in Cas9^+^ CD4^+^ T cells were performed 24 hours after T cell activation (0.25 μg/mL anti-CD3ε, 1 μg/mL anti-CD28) by centrifugation at 2000 rpm for 90 min at 32 ^o^C. Live and Thy1.1^+^ transduced cells cultured in Th17 polarizing conditions were analyzed by flow cytometry at 72 hrs.

### scRNA-seq and analysis

Colonic lamina propria cells from DSS or non-DSS treated mice were collected and enriched for CD4^+^ T cells using the mouse CD4^+^ T cell Isolation Kit (Miltenyi). Enriched CD4^+^ cells (~10,000 per mouse) were prepared for single cell libraries using the Chromium Single Cell 3’ Reagent Kit (10xGenomics). The pooled libraries of each sample (20,000 reads/cell) were sequenced on one lane of NovaSeq S4 following manufacturer’s recommendations. Cellranger v3.1.0 was used to filter, align, and count reads mapped to the mouse reference genome (mm10-3.0.0). The Unique Molecular Identifiers (UMI) count matrix obtained was used for downstream analysis using Seurat (v4.0.1). The cells with mitochondrial counts >5%, as well as outlier cells with a total gene number less than 200 were excluded. After filtering, batch effects in the samples with two biological repeats in DSS and non-DSS treated mice were evaluated and controlled by Seurat before downstream analysis. Initial cell clusters identified by Seurat with low expression of *Cd4* or *Cd3e* (mean log-transformed gene expression < 1) were removed, resulting in 10,308 total cells from eight samples. These cells were scaled and normalized using log-transformation, and the top 6,000 genes were selected for principal component analysis. The dimensions determined from PCA were used for clustering and UMAP visualizations.

Average gene expression in CPM (counts per million) in each cell clusters were used in calculation of fold changes between DSS and non-DSS treated samples. To avoid unconfident noise of gene expression occasionally observed in cell clusters, significant changes of gene expression in each cell cluster were identified here by a cutoff of 1.3-fold changes with averaged expression of each comparing pairs no less than 1 CPM in the DSS and non-DSS treated samples.

### RNA-seq

Ribosome-depleted RNAs were used to prepare sequencing libraries. 100 bp paired-end sequencing was performed on an Illumina HiSeq4000 by the Institute of Genomic Medicine at the University of California San Diego. Each sample yielded approximately 30-40 million reads. Paired-end reads were aligned to the mouse mm10 genome with the STAR aligner version 2.6.1a (71) using the parameters: “--outFilterMultimapNmax 20 --alignSJoverhangMin 8 -- alignSJDBoverhangMin 1 --outFilterMismatchNmax 999 --outFilterMismatchNoverReadLmax 0.04 --alignIntronMin 20 --alignIntronMax 1000000 --alignMatesGapMax 1000000”. Uniquely mapped reads overlapping with exons were counted using featureCounts (72) for each gene in the GENCODE.vM19 annotation. Differential expression analysis was performed using DESeq2 (v1.18.1 package) (73), including a covariate in the design matrix to account for differences in harvest batch/time points. Regularized logarithm (rlog) transformation of the read counts of each gene was carried out using DESeq2. Pathway analysis was performed on differentially expressed protein coding genes with minimal counts of 10, log_2_ fold change cutoffs of ≥ 0.4 or ≤-0.4, and p-values <0.05 using GeneCodis4.0.

### GRID-seq

GRID-seq of naïve and cultured Th17 cells were performed as described (51, 74). Briefly, two independent biological replicates (5-10 x10^6^) were crosslinked with disuccinimidyl glutarate (DSG) and formaldehyde. DNA in isolated nuclei were digested with AluI. A biotinylated bivalent linker was ligated to chromatin-associated RNA (with the ssRNA stretch on the linker) and nearby fragmented genomic DNA and captured by streptavidin microbeads for library construction. Single-end sequencing was performed on HiSeq400 (Illumina, 200 million reads/sample). Raw sequencing reads were evaluated by FastQC (75). Reads below 85bp were filtered out and those above 90bp with high quality were trimmed by Cutadapt (76) as suggested according to previous report (51, 74). The GRID-seq linker position at each read and the paired reads originated from RNA or genomic DNA were identified by matefq in GridTools (74). Paired RNA and DNA reads were then mapped to the mouse genome (GRCm38/mm10) by BWA respectively. Uniquely paired reads were used to generate a set of RNA-DNA interaction matrix for downstream analyses in the GridTools pipeline. Read counts from the two repeats were summarized into two 1kb genomic bins. Chromatin enriched with Malat1 RNAs (GRID-seq peak call) were defined as 2kb regions with clustered Malat1 signals above the background signal expected from random interactions (> 5-fold changes). In total, 7,039 regions out of 30,321 bins were identified as enriched Malat1 peaks for further analysis. Interactions of Malat1 RNA with Th17 genomic DNA were plotted in circos by circlize (75).

### ChIP-qPCR, ChIP-seq, and ATAC-seq

ChIP-qPCR and ChIP-seq were performed on 5-10 million crosslinked Th17 cells. For EZH2 and SUZ12 ChIP-seq assays, cells were cross-linked with 2 mM disuccinimidyl glutarate (DSG, ProteoChem) and 1% (vol/vol) formaldehyde (Thermo Fisher Scientific). H3K27me3 and H3K27ac were crossed with 1% formaldehyde only. Crosslinked cells were resuspended in ice-cold hypotonic buffer (10 mM HEPES-KOH (pH 7.9), 85 mM KCl, 1 mM EDTA, 1% NP-40, 1 mM PMSF, 0.5 mM sodium butyrate (Sigma-Aldrich), 1X protease inhibitor cocktail), and centrifuged at 1000 g for 5 min at 4°C to obtain a nuclear fraction. Nuclear pellets were resuspended in either LB3 buffer for H3K27me3 ChIP-seq (10 mM Tris-HCl (pH 7.5), 100 mM NaCl, 1 mM EDTA, 0.5 mM EGTA, 0.1% sodium deoxycholate, 0.5% N-laurosylsarcosine, 1 mM PMSF, 0.5 mM sodium butyrate, 1X protease inhibitor cocktail) or PIPA-NR buffer for SUZ12 ChIP-seq (20 mM Tris-HCl (pH 7.5), 150 mM NaCl, 1 mM EDTA, 0.5 mM EGTA, 0.4% sodium deoxycholate, 0.1% SDS, 1% NP-40, 0.5 mM DTT, 1 mM PMSF, 0.5 mM sodium butyrate, 1X protease inhibitor cocktail). Chromatin DNA was sonicated using a Covaris E220 and immunoprecipitated with antibodies pre-bound Dynabeads A/G overnight at 4°C. Immunoprecipitated protein-DNA complexes were reverse cross-linked and chromatin DNA purified to generate libraries as described in (77). The amplified libraries were purified, eluted and size selected using PAGE/TBE gel for 225-325 bp fragments by gel extraction, and single-end sequenced on NovaSeq 6000 (Illumina). ChIP-qPCR primer sequences are listed in Table S3.

ATAC-seq libraries were generated as described in (78). ChIP-seq and ATAC-seq processing followed the ENCODE guideline with some modifications (79). Specifically, single-end raw reads were mapped to the mouse genome (GENCODE assembly mm10) by bowtie2 (Version 2.3.4.1) in the local mapping mode with parameter “--local”, followed by PCR deduplication by SAMTools (Version 1.9) with the utility markedup (80). Mapped reads from each sample repeats were merged into a single BAM file by SAMTools, and peaks were called using MACS2 (Version 2.2.6) (81) in the narrow peak-calling mode with default parameters for ChIP-seq data or specific parameters of “callpeak --nomodel --extsize 100” for ATAC-seq data. Regions with peak-score below 30 were filtered out and the remaining reliable peak profiles were transformed into bigwig format and visualized on the Integrative Genomics Viewer (IGV Version 2.8.2) (82).

Super-enhancers were identified by normalized H3K27ac signals in genomic regions within a 12.5 Kb window outside of any known-gene promoters (TSS ± 2.5 kb) based on the algorithm as previously reported (16, 83). Briefly, any genomic regions calculated with overall H3K27ac coverage higher than the geometrically defined inflection point were considered putative superenhancers and those below were considered putative typical enhancers.

### cDNA synthesis and qRT-PCR

Total RNA was extracted with the RNeasy kit (QIAGEN) and reverse transcribed using Superscript III (Life technology). Real time RT-PCR was performed using iTaq™ Universal SYBR® Green Supermix (Bio-Rad Laboratories). Gene expressions were normalized to *Gapdh* mRNA levels. qRT-PCR primers were designed using Primer-BLAST to span across splice junctions, resulting in PCR amplicons that span at least one intron. Primer sequences are listed in Table S3.

### RNA FISH

RNA FISH was performed followed the protocol from BIOSERARCH TECHNOLOGIES. Briefly, cells were fixed with 3.7% formaldehyde and permeabilized with 70% ethanol, followed with hybridization with stellaris RNA FISH probes (Malat1-Cy3 and Il17f-Cy5). Nuclear DNA was visualized by 4’,6-diamidino-2-phenylindole (DAPI) and imaged with Leica SP8 confocal microscope.

### Statistics

All values are presented as means ± SD. Significant differences were evaluated using GraphPad Prism 8 software. The student’s t-test or paired t-test were used to determine significant differences between two groups. A two-tailed p-value of <0.05 was considered statistically significant in all experiments.

## Acknowledgments

We thank David Spector at Cold Spring Harbor Laboratory for sharing the Malat1^-/-^ mice. We thank Johannes Zuber and Aichinger Martin at the Research Institute of Molecular Pathology (I.M.P.) for sharing the pSIN vector for generating retroviruses to introduce custom sgRNAs into T cells.

S.M., N.C., C.L, Y.L., P.R.P, A.Z., and W.J.M.H. were partially funded by the National Institutes of Health (NIH) (R01 GM124494 to WJM Huang). B.Z. were partially funded by the Strategic Priority Research Program of the Chinese Academy of Sciences (XDA16010113), the National Key Research and Development Program of China (2019YFA0110002 and 2019YFA0801700) and the CAS Hundred Talents Program. Y.A. was funded by Japan Society for the Promotion of Science Overseas Research Fellowship 201860150, Uehara Memorial Foundation Fellowship, and NIH (R01 DK091183). S Patel and J Chang were partially funded by NIH (P01AI132122, and P30DK120515) and Biomedical Laboratory Research & Development Service of the VA Office of Research and Development (BX005106). Y.H. was supported by NIH grants HG004659 and DK098808. Illumina sequencing was conducted at the IGM Genomics Center, University of California San Diego, with support from NIH (S10 OD026929). The Moores Cancer Center Histology Core conducted colonic tissue sectioning and staining with support from NIH (P30 CA23100).

## Supplement Figures

**Fig S1.**
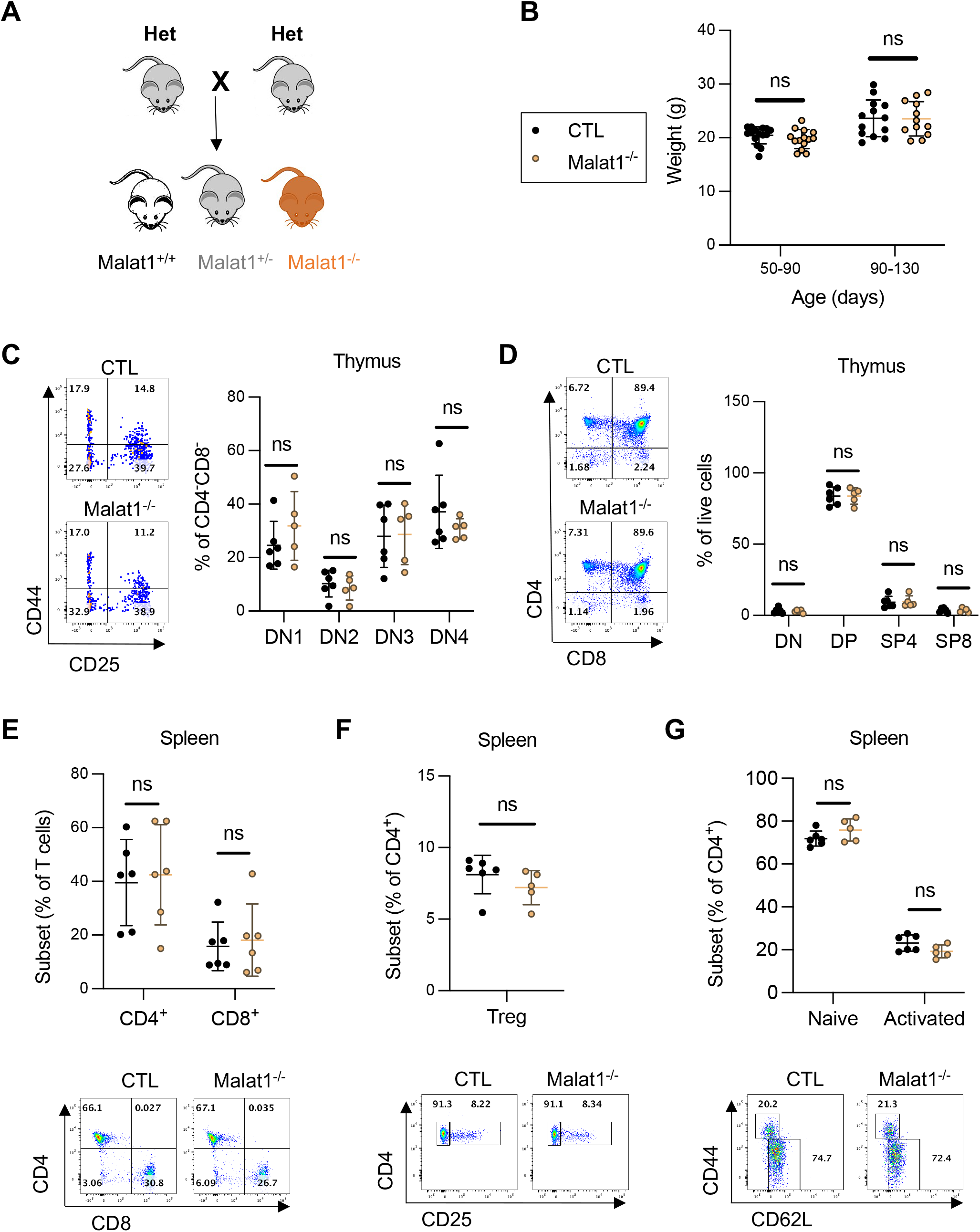
Normal thymic T cell development and splenic T cell populations in Malat1^-/-^ mice. A. Heterozygous crosses to generate Malat1^+/+^, Malat1^+/-^, and Malat1^-/-^ gender matched and cohoused littermates for experiments described in this study. B. Weight of CTL (Malat1^+/+^ or Malat1^+/-^) (n=30) and Malat1^-/-^ (n=28) mice assessed at the indicated ages. Each dot represents result from one mouse. ns: no significant (t-test). C. Representative flow cytometry analysis (left) and summarized percentage (right) of thymic DN subsets (DN1: CD44^-^CD25^-^, DN2: CD44^+^CD25^-^, DN3: CD44^+^CD25^+^, and DN4: CD44^-^ CD25^+^) from CTL and Malat1^-/-^ littermates. ns: not significant (multiple paired t-test, n=5). D. Representative flow cytometry analysis (left) and summarized percentage (right) of thymic developing T cells (DN: CD4^-^CD8^-^, SP4: CD4^+^CD8^-^, SP8: CD4^-^CD8^+^, and DP: CD4^+^CD8^+^) from CTL and Malat1^-/-^ littermates. ns: not significant (multiple paired t-test, n=6). E. Summarized percentage (top) and representative flow cytometry results (bottom) of CD4 and CD8 expression in total splenocytes from CTL and Malat1^-/-^ littermates. ns: not significant (multiple paired t-test, n=6). F. Summarized percentage (top) and representative flow cytometry results (bottom) of Tregs (CD25^+^) among splenic CD4^+^ cells from CTL and Malat1^-/-^ littermates. ns: not significant (paired t-test, n=6). G. Summarized percentage (top) and representative flow cytometry results (bottom) of splenic naïve (CD44^-^CD62L^+^) and activated (CD44^+^CD62L^-^) CD4^+^ T helper cells from CTL and Malat1^-/-^ littermates. ns: not significant (multiple paired t-test, n=5).

**Fig S2.**
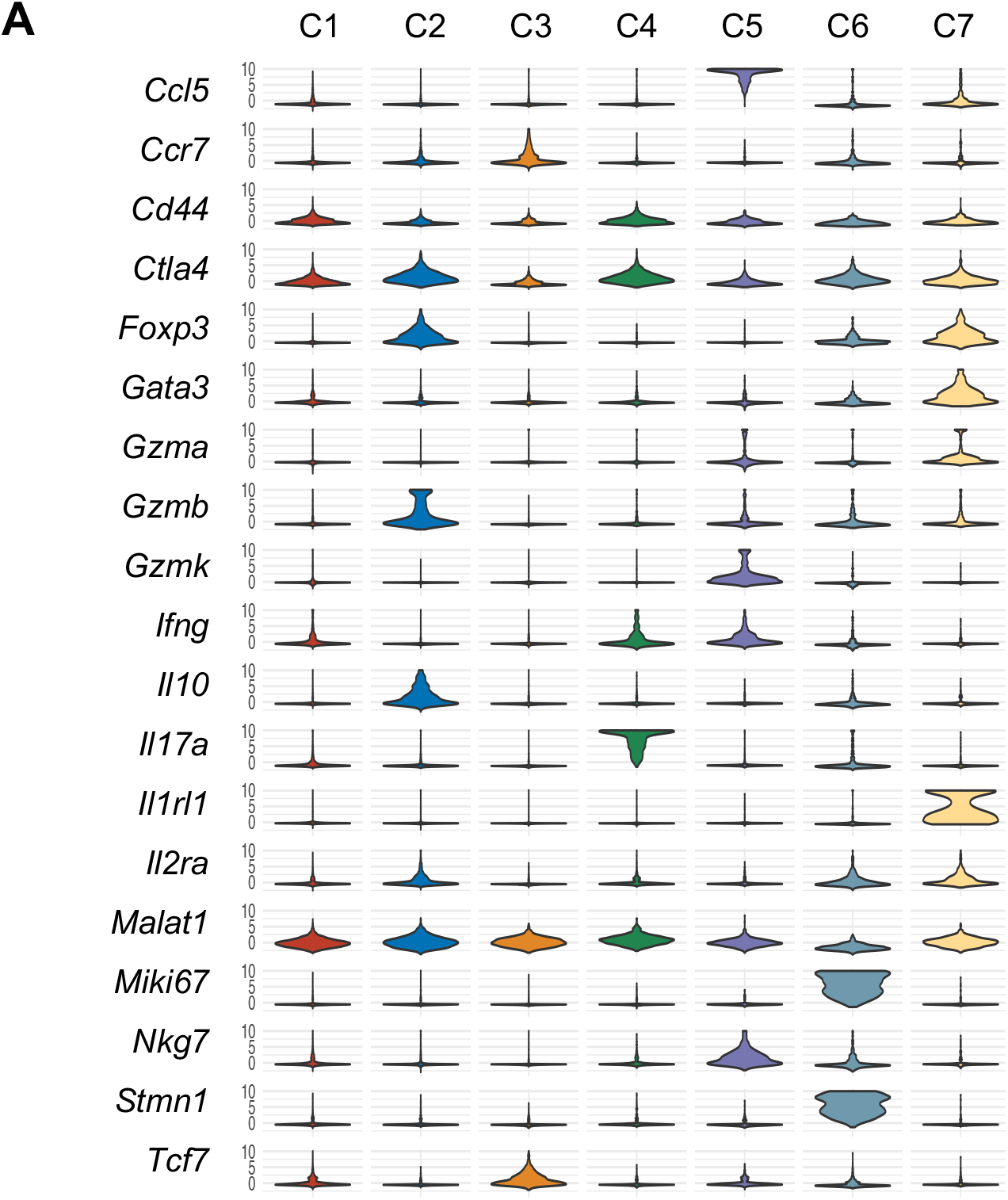
Select gene expression in different CD4^+^ T cell clusters. A. Expression of select cluster-specific genes in colonic CD4^+^ T cell clusters as determined by scRNA-seq.

**Fig S3.**
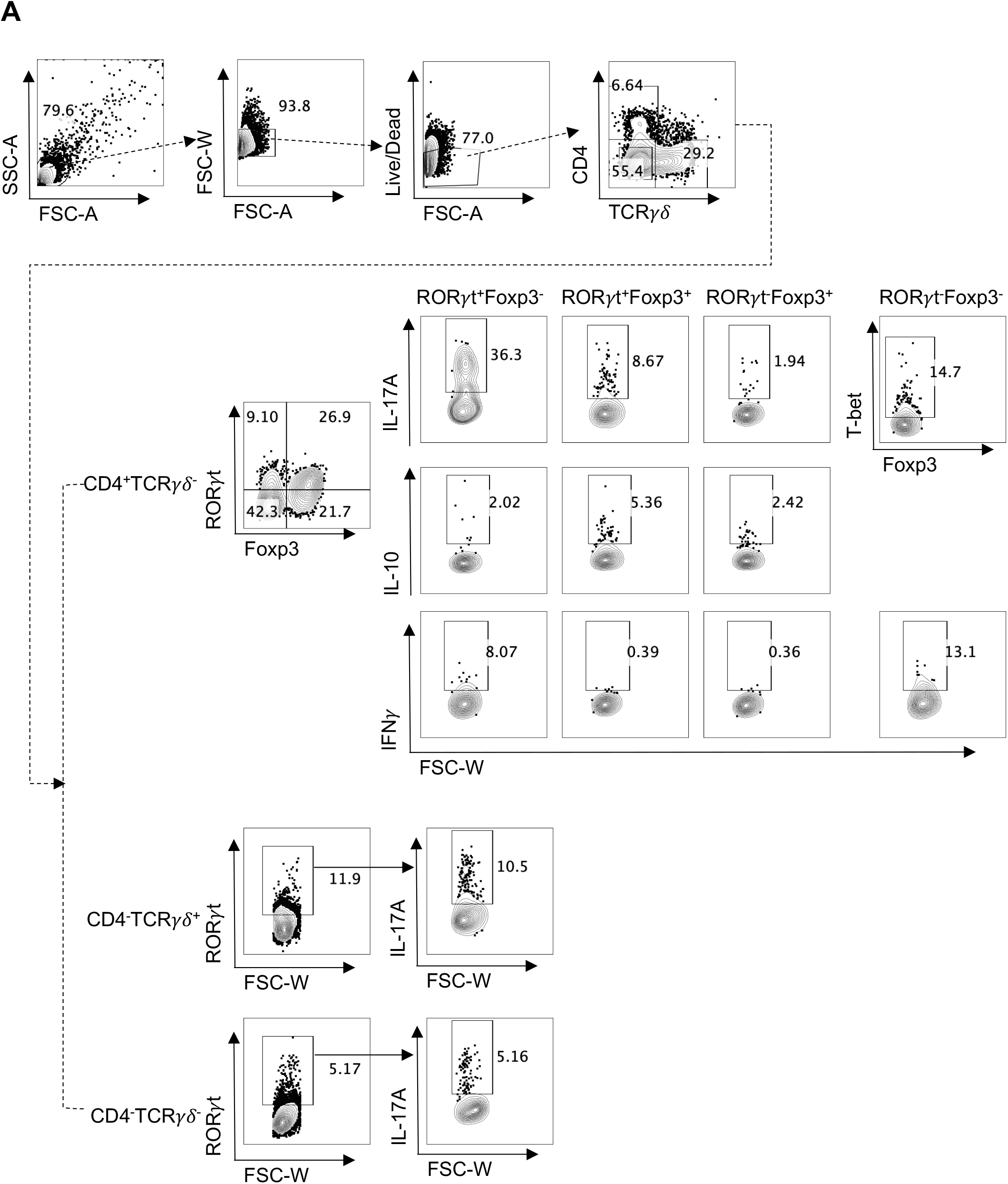
Gating strategy of immune T cell in colonic lamina propria. A. Cells were gated for lymphocytes by FSC and SSC, doublets were discriminated, and live cells were analyzed for CD4 and TCRγδ expression. CD4^+^ T cells were further analyzed for subpopulations according to RORγt and/or Foxp3 expression, and cytokines expressions were further analyzed in those subpopulations.

**Fig S4.**
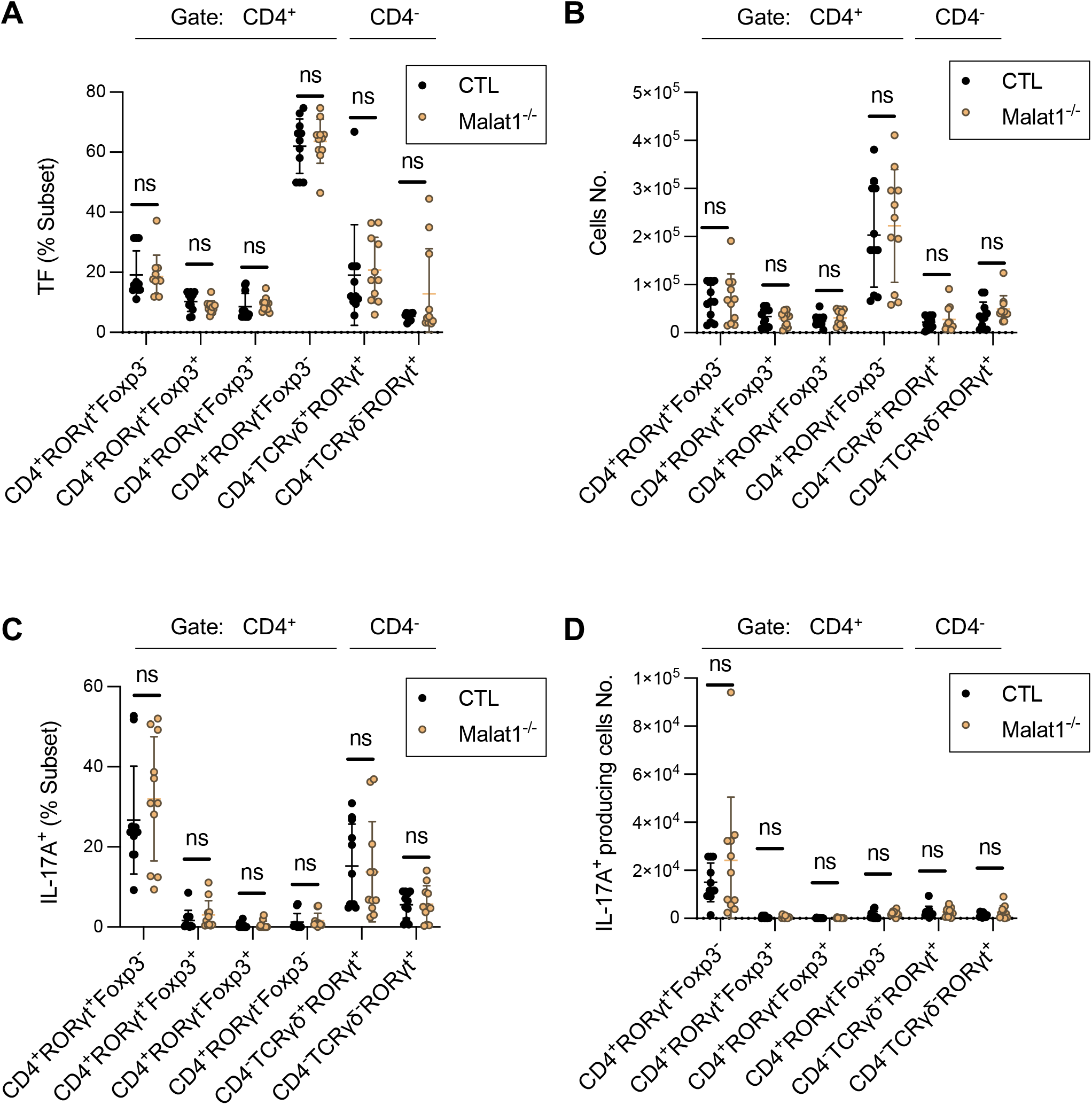
Colonic T cell populations in CTL and Malat1^-/-^ mice under steady state. A. Proportion of indicated cells in steady state cLP of CTL (n=11) and Malat1^-/-^ (n=11) mice. Each dot represents result from one mouse. ns: not significant (multiple t-test). B. Cell number of A. C. Proportion of IL-17A in indicated steady state cLP of CTL (n=11) and Malat1^-/-^ (n=11) mice. Each dot represents result from one mouse. ns: not significant (multiple t-test). D. Cell number of C.

**Fig S5.**
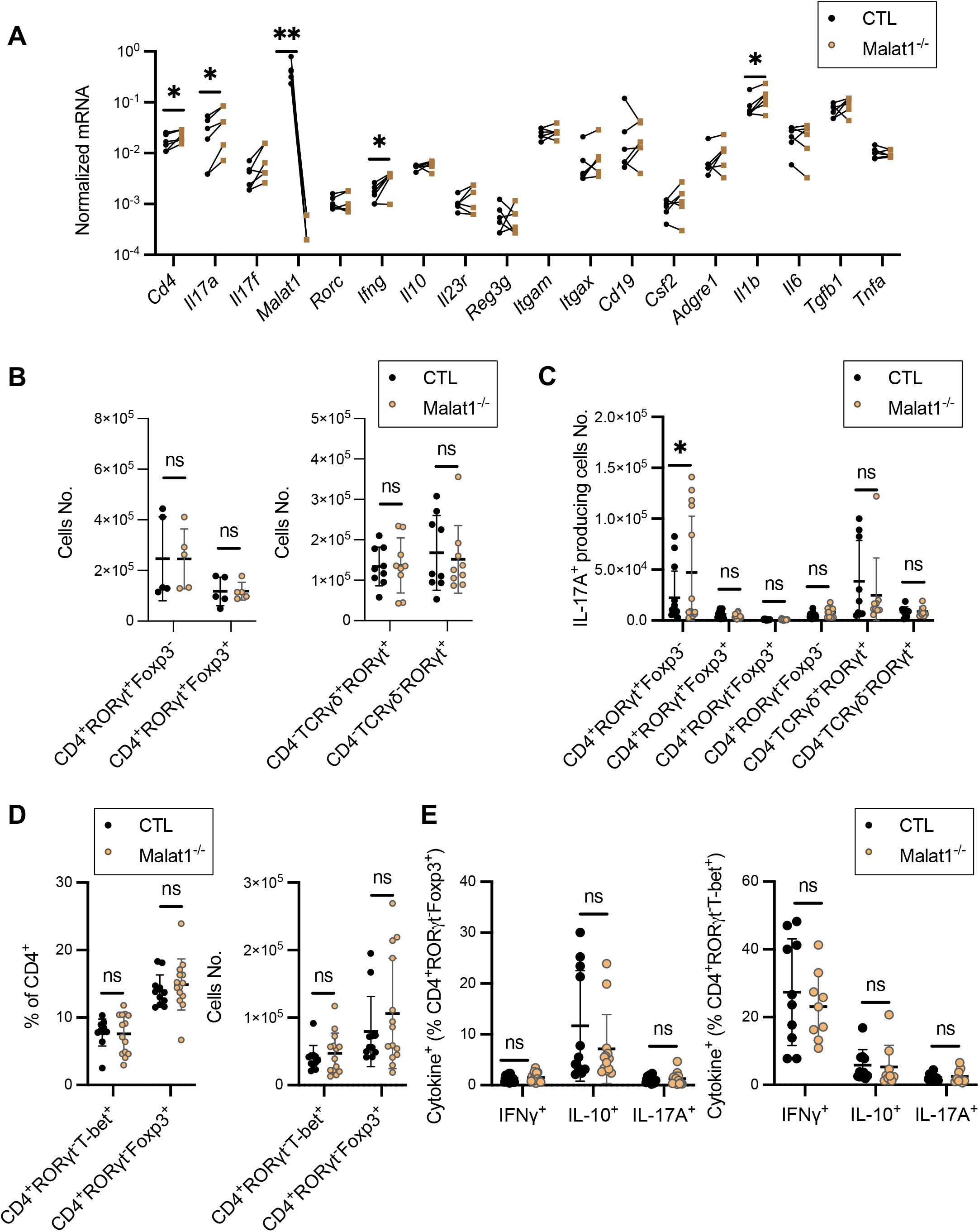
Cytokine expressions in different colonic T cell populations post DSS treatment. A. Relative mRNA level of the indicated genes in total colonic lamina propria mononuclear cells harvested from CTL (n=6) and Malat1^-/-^ (n=6) mice on day 10 post initial DSS treatment as detected by qRT-PCR. Each dot represents result from one mouse. * p-value<0.05, ** p-value<0.01, ns: not significant (multiple paired t-test). B. Left: cell number of cLP Th17 (CD4^+^ RORγt^+^Foxp3^-^) and RORγt^+^ Treg (CD4^+^RORγt^+^Foxp3^+^) from CTL (n=5) and Malat1^-/-^ (n=5) littermates harvested on day 10 post DSS treatment. Right: cell number of cLP Tγδ17 (CD4^-^TCRγδ^+^RORγt^+^) and ILC3 (CD4^-^TCRγδ^-^RORγt^+^) from CTL (n=9) and Malat1^-/-^ (n=9) littermates harvested on day 10 post DSS treatment. ns: not significant (multiple paired t-test). C. Cell number of indicated IL-17A producing cells in cLP from CTL and Malat1^-/-^ mice harvested on day 10 post DSS treatment. Each dot represents result from one mouse (multiple t-test). D. Proportion (left) and cell number (right) of indicated CD4^+^ T cells in cLP from CTL and Malat1^-/-^ mice harvested on day 10 post DSS treatment. Each dot represents result from one mouse. ns: not significant (multiple t-test). E. The proportion of cLP Treg and Th1 cells expressing the indicated cytokines from the of DSS treated CTL and Malat1^-/-^ mice harvested on day 10. Each dot represents result from one mouse. ns: not significant (multiple t-test).

**Fig S6.**
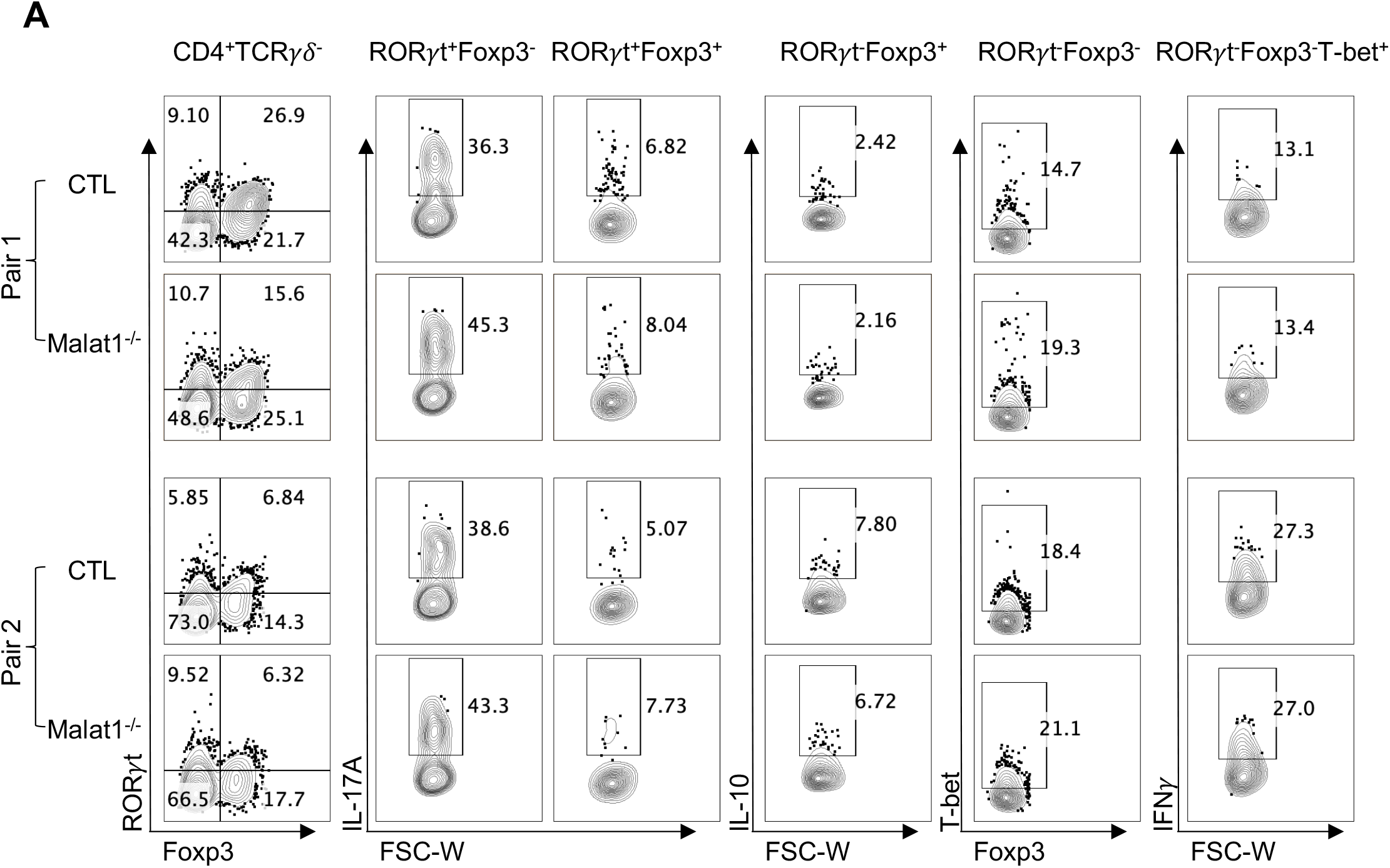
Representative flow analysis of colonic CD4^+^ T cell subsets. A. Representative flow analysis of CD4^+^ subsets and their cytokines potential in two pairs of from CTL and Malat1^-/-^ littermates harvested on day 10 post DSS treatment.

**Fig S7.**
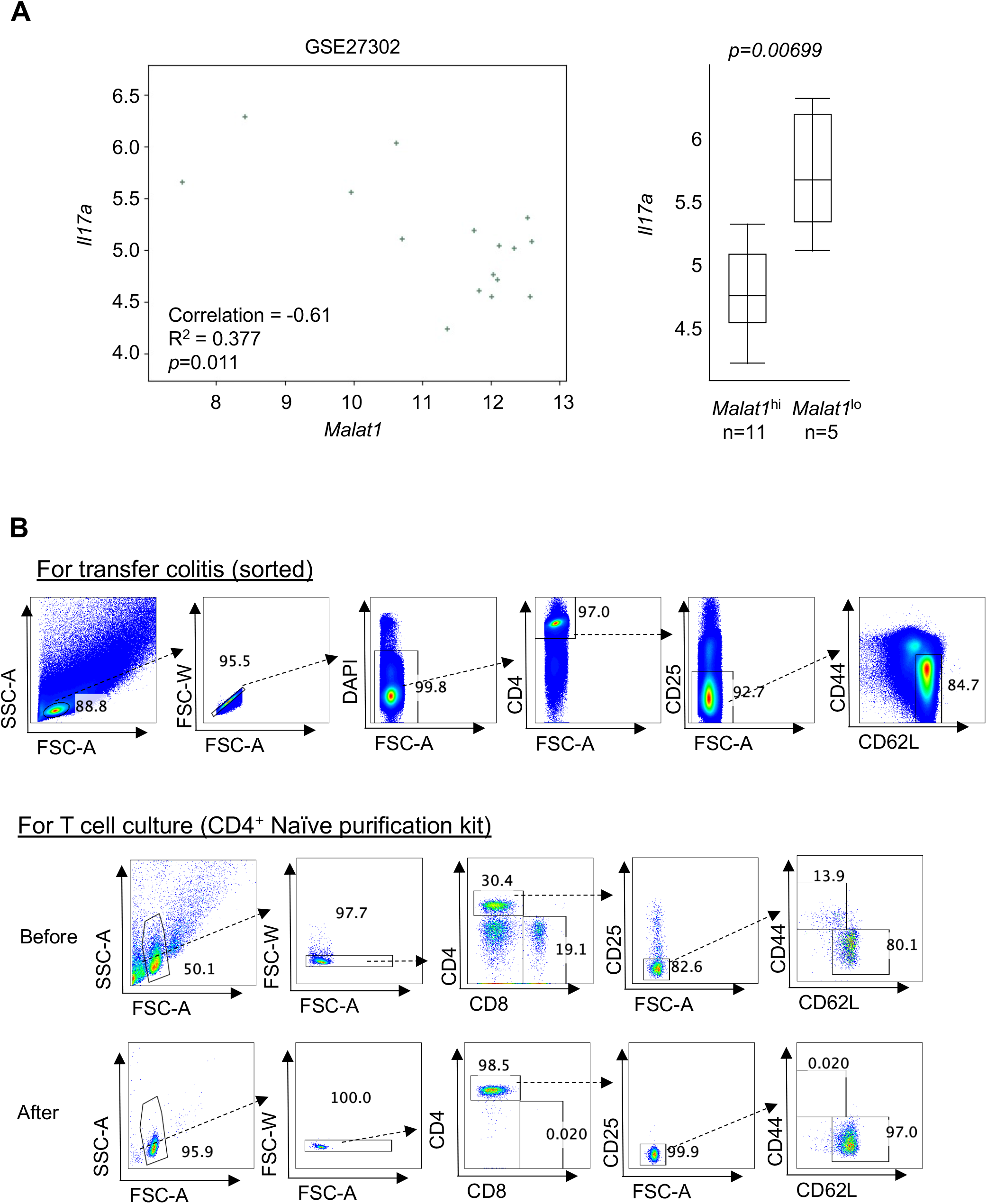
Select colonic gene expression in the CD4^+^ T cell transfer colitis model. A. Scatter plot of RNA level of *Malat1* and *Il17a* in colonic tissues from the CD4^+^ T cell transfer colitis model (GSE27302, (48)). B. Naïve CD4^+^ T cell sorting strategy (top). Confirmation of naïve CD4^+^ T cell purity post isolation using the naïve CD4^+^ T cell isolation kit (Miltenyi; bottom).

**Fig S8.**
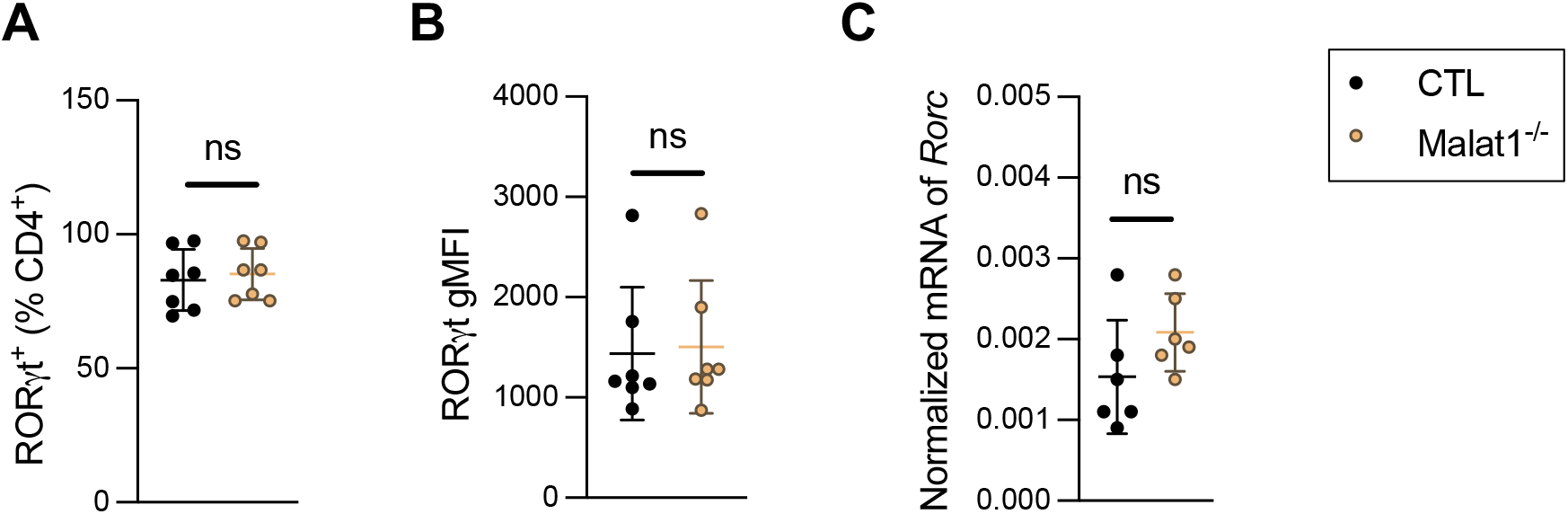
RORγt protein expression in cultured Th17 cells. A. Summarized proportion of RORγt^+^ Th17 cells from CTL and Malat1^-/-^ littermates cultured with IL-6 and TGFβ (n=8). Each dot represents result from one independent culture. ns: not significant (paired t-test). B. Summarized RORγ gMFI of cells described in A. ns: not significant (paired t-test). C. Summarized mRNA level of *Rorc* in cells described in A. ns: not significant (paired t-test).

**Fig S9.**
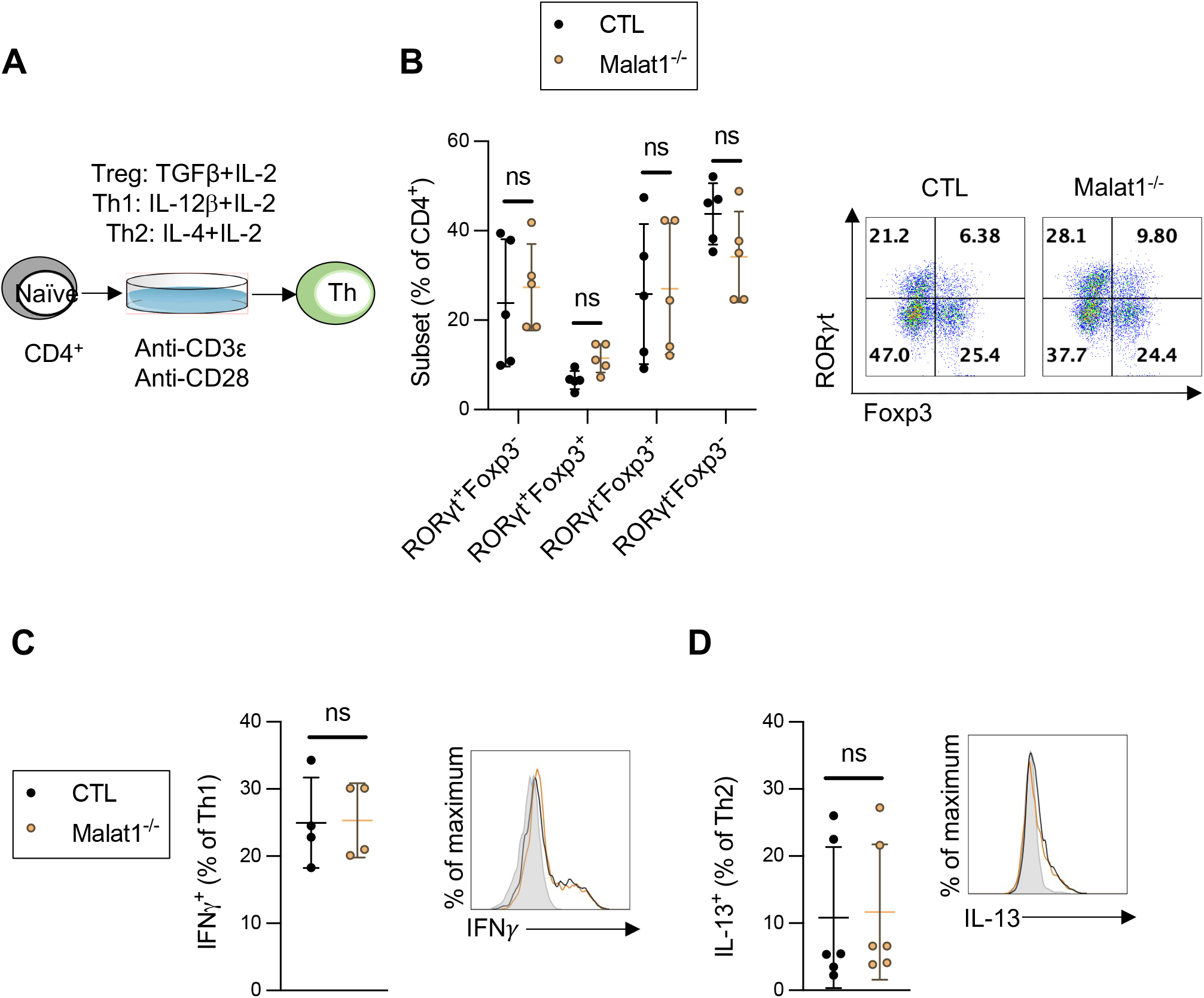
Malat1 does not regulate Th1, Th2, and Treg generation and function *in vitro*. A. Workflow for polarizing naïve CD4^+^ T cell toward different T cell subsets *in vitro*. B. Summarized proportion (left) and representative flow cytometry analysis (right) of RORγt and/or Foxp3 expressing T cells from CTL (n=5) and Malat1^-/-^ (n=5) littermates cultured in the presence of TGFβ and IL-2. Each dot represents result from one independent culture. ns: not significant (multiple paired t-test). C. Summarized proportion (left) and histogram (right) of IFNγ expression in cultured Th1 cells from CTL (n=4) and Malat1^-/-^ (n=4) littermates. ns: not significant (paired t-test). D. Summarized proportion (left) and histogram (right) of IL-13 expression in cultured Th2 cells from CTL (n=6) and Malat1^-/-^ (n=6) littermates. ns: not significant (paired t-test).

**Fig S10.**
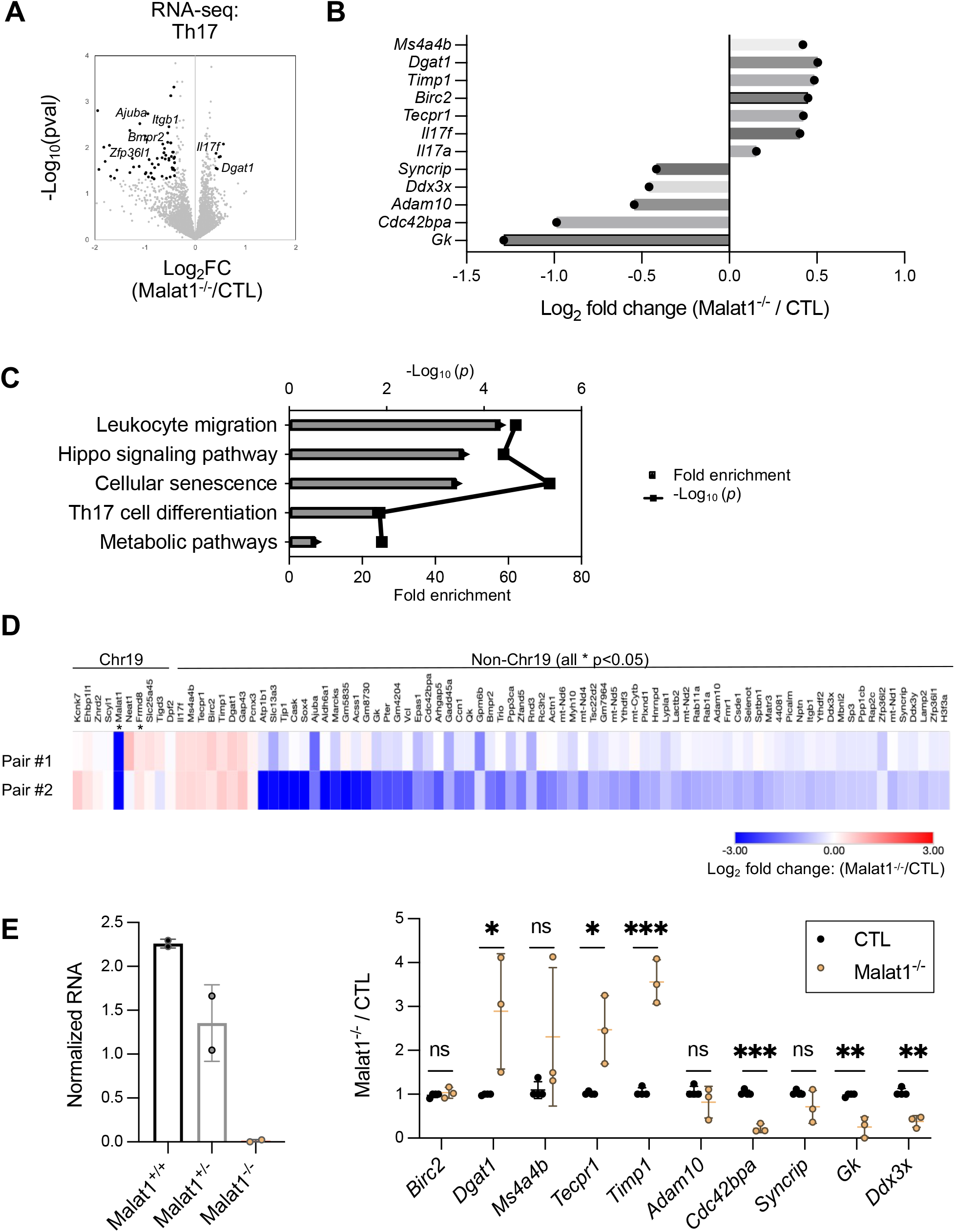
Malat1-dependent gene programs in cultured Th17 cells. A. Volcano plot depicting differential gene expression in Malat1^-/-^ and CTL Th17 cells generated in the presence of anti-CD3e, anti-CD28, IL-6, and TGFβ for 3 days from two independent experiments. Black dots: genes with a log_2_ fold-change > 0.4 or < −0.4, p-value < 0.05 and minimal counts of 100 (DEseq2) are highlighted in black. B. Log_2_ fold change of select Malat1-dependent genes in Th17 cells from A. C. Top KEGG pathways enriched with Malat1-dependent genes in Th17 cells from A. D. Heatmap displaying average log_2_ fold change of *Malat1* neighboring genes (chr19) and Malat1 dependent genes outside of chr19 in cultured CTL and Malat1^-/-^ Th17 cells. E. Left: normalized RNA expression of *Malat1* in cultured Th17 cells in *Malat1*^+/+^, *Malat1*^+/-^, and *Malat1^-/-^* mice. Right: normalized mRNA expression of the indicated genes in cultured Th17 cells from CTL and Malat1^-/-^ mice as detected by qRT-PCR. * p-value<0.05, ** p-value<0.01, *** p-value<0.001, ns: not significant (multiple paired t-test, n=5).

**Fig S11.**
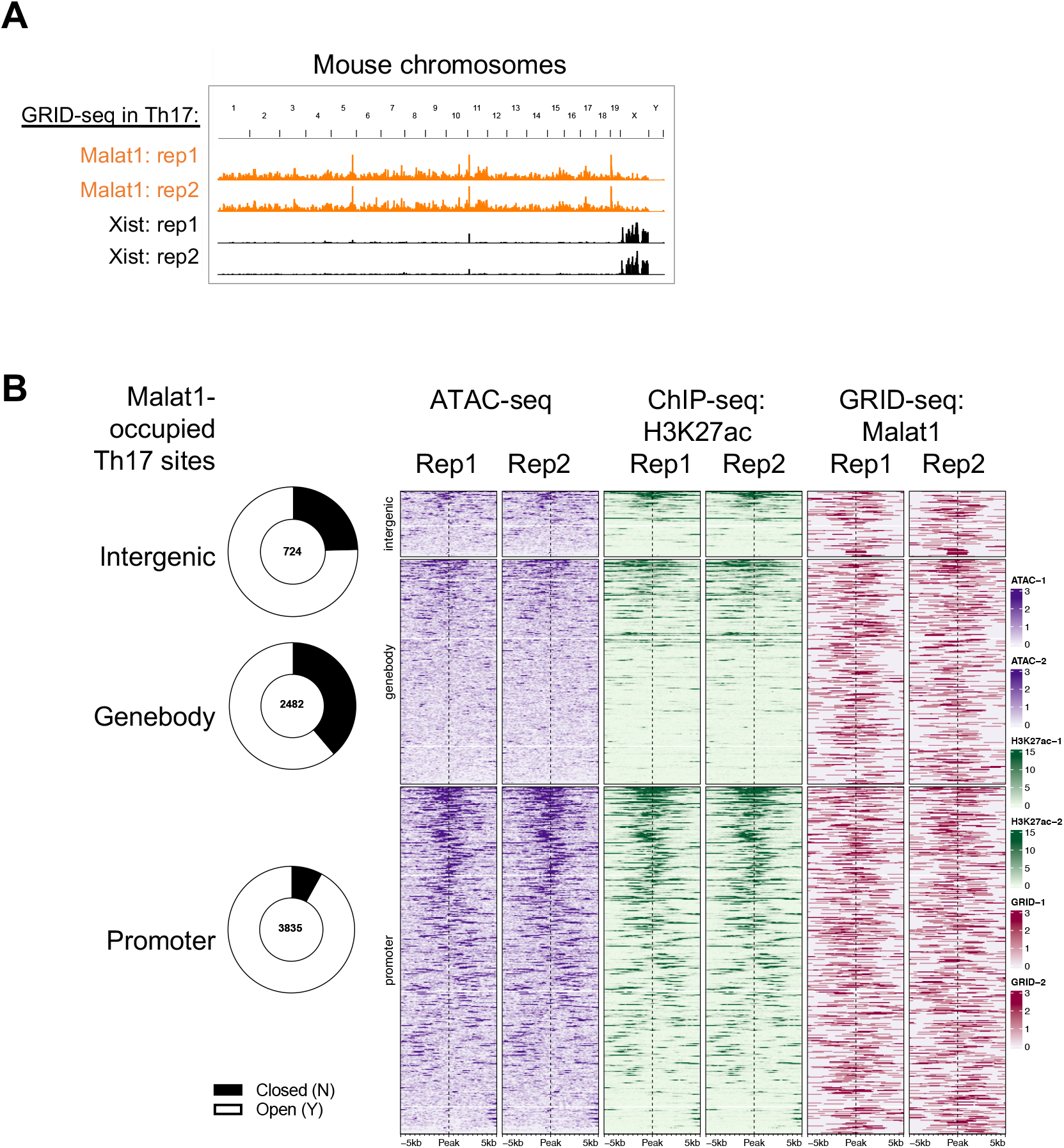
GRID-seq identified chromatin occupancy of lncRNAs in Th17 cells. A. GRID-seq signals of Malat1 and Xist displaying their genome-wide occupancy on the chromatin in two biological replicates of Th17 cells. B. Open chromatin (ATAC-seq), H3K27ac (ChIP-seq), and Malat1 (GRID-seq) signals on Malat1 occupying peaks. Th17 promoters (3,834 total) were defined as 2.5kb genomic regions flanking TSS; DREs (2,476 total) were defined as those within 12.5kb of a H3K27ac peak; gene bodies (6,166 total) were scaled displayed of first TSS to the last TTS of each annotated gene.

**Fig S12.**
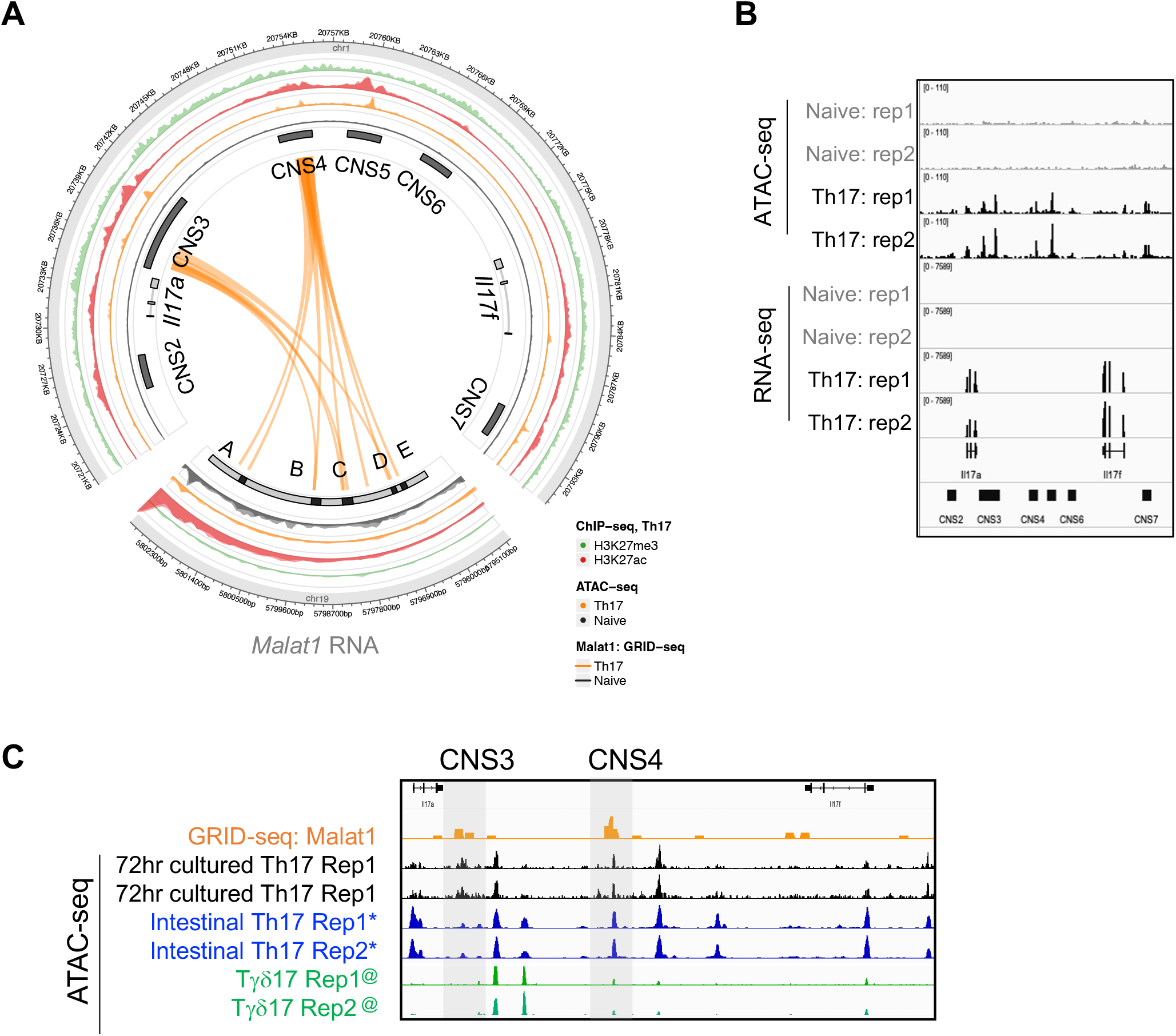
Malat1 occupies select regulatory elements at the *Il17a-Il17f* super-enhancer in cultured Th17 cells. A. Circos plot showing Malat1 trans-interactions (Th17: orange lines, Naïve: grey line) detected by GRID-seq at the *Il17a-Il17f* locus in CTL Th17 cells. Five Malat1 RNA regions enriched on the *Il17a-Il17f* locus were designated as A-E. The outer circles show signals of ATAC-seq (grey: signals from two independent naive cell replicates; orange: signals from two independent Th17 replicates), H3K27ac (red: ChIP-seq signals from two independent Th17 replicates), and H3K27me3 (green: ChIP-seq signals from GSE40918 (85)). B. IGV browser of ATAC-seq and RNA-seq results from two independent replicates of wildtype naïve and Th17 cells at the *Il17a-Il17f* locus. C. IGV browser showing ATAC-seq signal of the *Il17a-Il17f* locus in the indicated cells, including cultured Th17, intestinal Th17 cells (GSE127768 (55)), and Tγδ17 cells (GSE120426, (56)).

**Fig S13.**
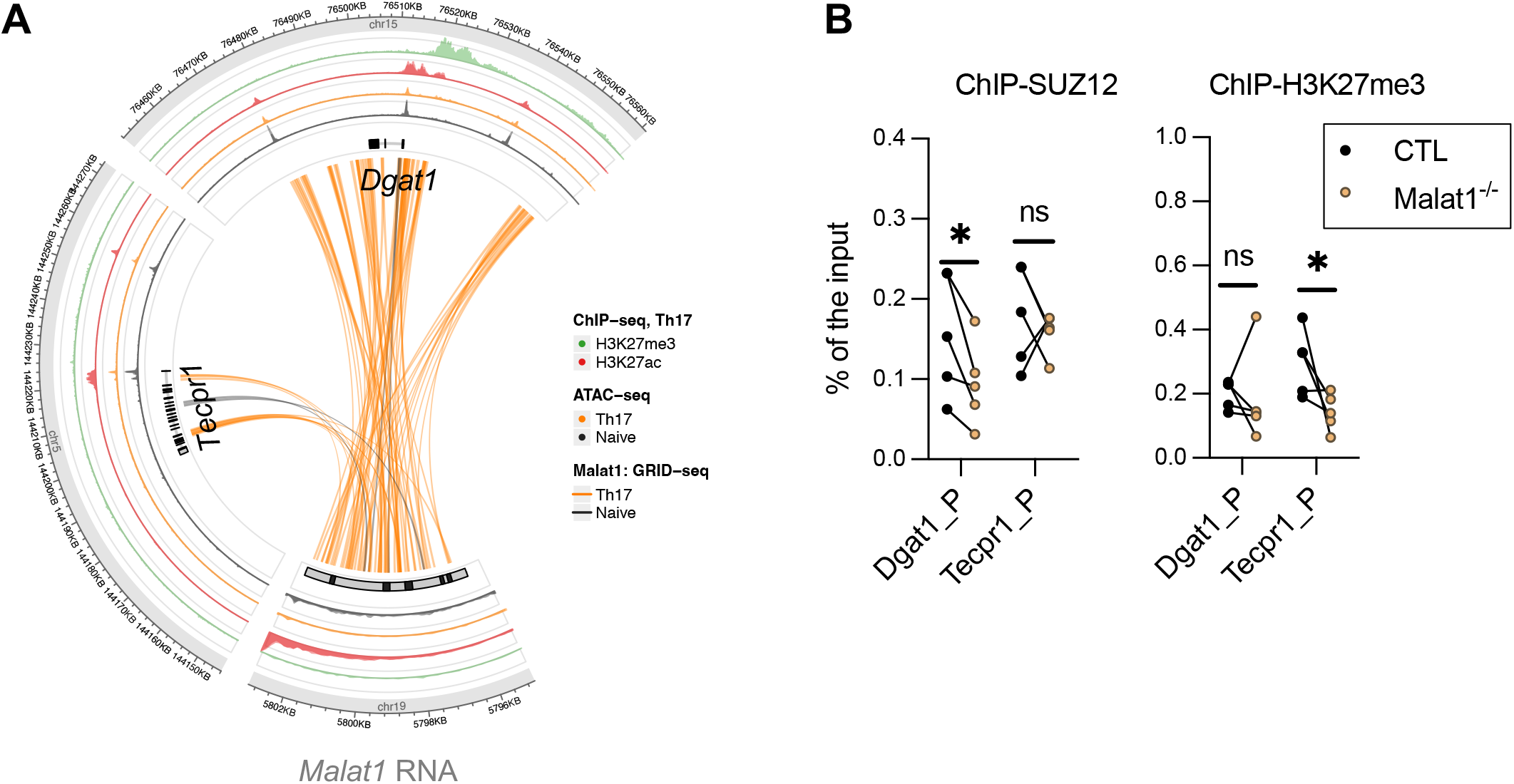
Malat1 chromatin occupancy on other direct target genes in Th17 cells. A. Circos plot showing Malat1 trans-interactions (Th17: orange lines, Naïve: grey line) detected by GRID-seq at the *Dgat1* and *Tecpr1* loci in CTL Th17 cells. The outer circles show signals of ATAC-seq (grey: signals from two independent naive cell replicates; orange: signals from two independent Th17 replicates), H3K27ac (red: ChIP-seq signals from two independent Th17 replicates), and H3K27me3 (green: ChIP-seq signals from GSE40918 (85)). B. Enrichment of SUZ12 and H3K27me3 on the promoters of *Dgat1* and *Tecpr1* as determined by ChIP-qRT. p, promoter. * p-value<0.05, ns: not significant (multiple paired t-test, n=5).

